# Antigen-presenting cancer-associated fibroblasts in murine pancreatic tumors differentially control regulatory T cell phenotype and function via CXCL9 and CCL22

**DOI:** 10.1101/2025.03.27.645833

**Authors:** Saumya Y. Maru, Meredith Wetzel, Jacob T. Mitchell, Nicole E. Gross, Lalitya Andaloori, Kathryn Howe, Emma Kartalia, Guanglan Mo, James Leatherman, Won Jin Ho, Elana J. Fertig, Luciane T. Kagohara, Edward J. Pearce, Elizabeth M. Jaffee

**Author notes:** **COI disclosure statement**: SYM reports honoraria from Omni Health Media. WJH reports patent royalties from Rodeo/Amgen, received research funding from Sanofi, NeoTX, Riboscience (to Johns Hopkins), and speaking/travel honoraria from Exelixis and Standard BioTools. EJF was on the SAB for Resistance Bio, received personal fees from Mestag and Merck, and grants from Roche/Genetech, Abbvie Inc. EJP is on the SAB for Remedy Plan Therapeutics. EMJ reports other support from Abmeta and Adventris, personal fees from Dragonfly, Neuvogen, CPRIT, Surge Tx, Mestag, Medical Home Group, HDTbio, and grants from Lustgarten, Genentech, BMS, NeoTx, and Break Through Cancer. EMJ is the Dana and Albert “Cubby” Broccoli Professor of Oncology.

## Abstract

Pancreatic ductal adenocarcinoma (PDAC) is characterized by a complex tumor microenvironment (TME) including stromal cells that influence resistance to therapy. Recent studies have revealed that stromal cancer-associated fibroblasts (CAFs) are heterogeneous in origin, gene expression, and function. Antigen-presenting CAFs (apCAFs), are defined by major histocompatibility complex (MHC)-II expression and can activate effector CD4^+^ T cells that have the potential to contribute to the anti-cancer immune response, but also can induce regulatory T cell (Treg) differentiation. Whether apCAFs promote or restrain the antitumor response remains uncertain. Using tumor clones of the KPC murine PDAC model differing in sensitivity to immune checkpoint blockade (ICB), we found that immunosensitive (sKPC) tumors were characterized by higher immune cell and apCAF infiltration than resistant (rKPC) tumors. IMC analysis showed proximity of apCAFs and CD4^+^ T cells in both sKPC and rKPC tumors implicating interaction within the TME. apCAF-depleted sKPC tumor-bearing mice had diminished sensitivity to ICB. apCAFs from both sKPC and rKPC tumors activated tumor-infiltrating CD4^+^ T cells and induced Treg differentiation. However, transcriptomic analysis showed that Tregs induced by apCAFs were overexpressed for immunosuppressive genes in rKPCs relative to sKPCs, and that this is associated with differential chemokine signaling from apCAFs depending on tumor origin. Together these data implicate apCAFs as important mediators of the antitumor immune response, modulation of which could facilitate the development of more effective anti-tumor immune based approaches for PDAC patients.

## Introduction

Pancreatic ductal adenocarcinoma (PDAC) is among the most aggressive malignancies with a five-year overall survival of only 11% (1). New drug development has been slow and the existing standard of care cytotoxic chemotherapy regimens are poorly tolerated and have limited clinical benefit. Outside of a small minority of PDAC patients with mismatch repair deficient (MMR-d) or microsatellite instability (MSI-h) tumors, immunotherapy does not confer significant antitumor activity (2). Even within MMR-d and MSI-h PDAC, rates of disease control and cure with immunotherapy are inferior to rates achieved in other GI malignancies such as colorectal carcinomas (3). Emerging data suggest that multiple cellular components especially stromal cells in the PDAC tumor microenvironment (TME) contribute to primary resistance to most therapies including immunotherapy (4,5). Understanding the role of these stromal cells and their regulation will facilitate the development of more effective anti-tumor immune based approaches for PDAC patients.

The PDAC TME is characterized by a high degree of desmoplasia and cancer-associated fibroblasts (CAFs) have been implicated as contributors to poor clinical outcomes (4,5). CAFs are a heterogeneous group in phenotype and function, consisting of populations such as myofibroblastic CAFs (myCAFs), inflammatory CAFs (iCAFs), and the more recently described subset, antigen-presenting CAFs (apCAFs) (4,6,7). apCAFs, defined by expression of MHC-II, can present antigens to CD4^+^ T cells and regulate CD4^+^ T cell phenotype (8,9). Lineage tracing studies found that apCAFs have a distinct origin from traditional fibroblasts and are derived from mesothelium rather than mesenchyme (8). Furthermore, spatial transcriptomics studies have revealed the presence of apCAFs near early precancerous lesions, pancreatic intraepithelial neoplasia (PanIN) (10). These features make apCAFs attractive as a unique cell subset and potential target for modulation within the TME; however, there are conflicting reports as to whether apCAFs promote or restrain the anti-tumor response. For example, an early study of apCAF function found antigen-specific induction of regulatory T cell (Treg) differentiation and decreased tumor weight following apCAF depletion in a T cell inflamed model of murine PDAC (8), implicating a tumor-promoting role of apCAFs. apCAFs have been associated with expression of SPP1, which facilitates both primary tumor formation and metastases (11,12). Conversely, several studies have suggested that immunofibroblasts, CAFs with inflammatory and interferon-response features, are associated with good patient prognosis and tumor restraint (13–15). Most recently, gastric cancer patients with tumors enriched for apCAFs were found to have increased overall survival (9).

In this study we isolated apCAF and CD4^+^ T cell interactions to better understand the impact of this cellular crosstalk on the antitumor response as a function of the TME. We used tumor clones derived from the autochthonous K-ras^LSL.G12D/+^; p53^R172H/+^; PdxCre (KPC) murine PDAC model (16,17) that differ in sensitivity to immune checkpoint blockade (ICB) to establish orthotopic pancreatic tumors and observed that apCAFs are more abundant in tumors arising from immunosensitive tumors, and increase in sensitive tumors following ICB. Furthermore, depleting apCAFs in tumor-bearing mice decreased the response of immunosensitive tumors to ICB. Imaging mass cytometry (IMC) studies showed that apCAFs are spatially involved with CD4^+^ T cells in both the sensitive and resistant TME and ex-vivo cocultures demonstrated that apCAFs from both environments activate CD4^+^ T cells and induce Treg differentiation. However, single-cell RNA sequencing (scRNAseq) experiments showed differential chemokine signaling from apCAFs depending on tumor of origin that indicated variable transcriptional programming within Tregs. apCAFs from immunoresistant tumors induced classical Tregs via CCL22 signaling and had a higher immunosuppressive gene signature, whereas apCAFs from the immunosensitive TME signaled via CXCL9. Together these findings implicate apCAFs as important mediators of the antitumor immune response in PDACs.

## Results

### Orthotopically implanted sKPC and rKPC tumors differ in responsiveness to immune checkpoint blockade (ICB)

We first validated two KPC tumor clones that have been previously characterized as T cell high or low to establish an *in-vivo* system allowing us to compare responsiveness to standard ICB (17). YFP-tagged KPC clones 2838c3 (T cell high) and 6419c5 (T cell low) were orthotopically implanted into the pancreas of syngeneic C57BL/6 mice as previously described (18,19). After allowing tumors to establish for 10 days we began twice weekly intraperitoneal injections of anti-PD1 and anti-CTLA4 antibodies or respective isotype controls and continued dosing until day 30 (**Figure 1A**). Mice were monitored for death or extremis endpoints as required by our institutional animal protocol. Survival curves (**Figure 1B-C, Table 1**) demonstrated marked difference in responsiveness to anti-PD1 and anti-CTLA4 treatment as has been previously described (17). 6419c5 tumors are more aggressive than 2838c3 tumors with a median survival of 21 days and 37 days respectively. 6419c5 tumor-bearing mice had minimal response to dual ICB, although in contrast to prior studies there was a significant difference in medial survival (21 versus 27 days), likely due to drift that occurs with repeated passaging of tumor cells. 2838c3 tumor-bearing mice, however, had a greater response to treatment, with median survival lengthening from 27 days for isotype-treated mice to 59.5 days for mice receiving dual ICB. At 120 days two out of ten ICI-treated mice initially implanted with 2838c3 tumors were alive with no evidence of tumor. Separately 2838c3 tumor-bearing mice were treated with dual ICB of isotype control as described above and sacrificed at day 28 for assessment of tumor weight. Mice treated with dual ICB had significantly smaller tumors (**Figure 1D**). We thus refer to 2838c3 tumors as sensitive (sKPC) and 6419c5 tumors as resistant (rKPC) to ICB. Together, these data validate the use of T cell low and T cell high KPC tumor clones for further exploration of the TME.

**Figure 1:**
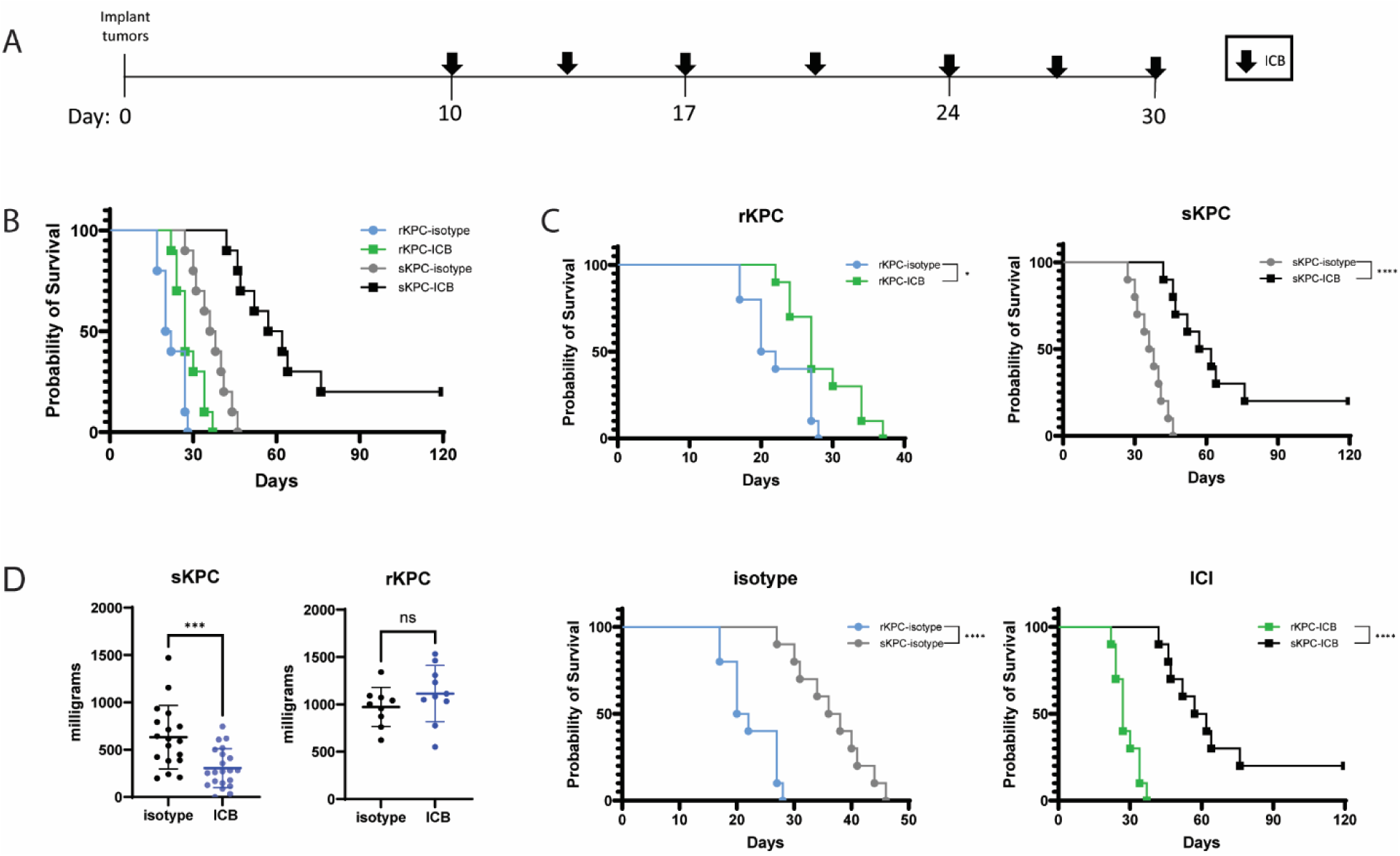
Orthotopically implanted sKPC and rKPC tumors differ in responsiveness to immune checkpoint blockade. **(A)** Experiment timeline. sKPC (KPCY clone 2838c3) and rKPC (KPCY clone 6419c5) tumors were implanted orthotopically at day 0. ICB was started on day 10 and continued twice weekly through day 30. Mice were monitored for signs of extremis in accordance with institutional guidelines. **(B-C)** Survival curves for orthotopic sKPC and rKPC tumor-bearing mice treated with ICB or isotype-control antibodies, n= 10 mice per group. Survival curves are shown for all four experimental conditions (B) and as pairwise comparisons of tumor type and treatment (C). **(D)** Tumor weights of sKPC and rKPC tumor-bearing mice ± ICB at day 28 post implantation.

**Table 1:** Median survival of sKPC and rKPC tumor-bearing mice ± ICB, n= 10 mice per group.

| Group | Median survival (days) |
| --- | --- |
| rKPC isotype | 21 |
| rKPC ICI | 27 |
| sKPC isotype | 37 |
| sKPC ICI | 59.5 |

### Immunosensitive PDAC TME is characterized by T cell and apCAF infiltration

We next analyzed the TME of sKPC and rKPC tumors to characterize their cellular compartments. Orthotopically implanted sKPC and rKPC tumors were harvested after three weeks and dissociated to a single cell suspension for quantification of immune cells by flow cytometry. sKPC tumors are T cell high as compared to rKPC tumors, as previously described (17) (**Supplementary Figure 1**). There was no significant difference in Treg numbers between tumors, but we did observe an increase in T helper type 1 (Th1) cells in rKPC tumors. Similarly, we observed more CD11c^+^ dendritic cells in rKPC tumors (**Supplementary Figure 1**). Further analysis of the stromal component of the TME showed no difference in absolute counts of total CAFs between sKPC and rKPC tumors (**Figure 2A**). However, apCAFs were more prevalent in sKPC tumors (**Figure 2B**). We also observed a trend towards increased apCAF numbers when sKPC tumor-bearing mice were treated with dual ICB (**Figure 2C**), suggesting that apCAF prevalence may also serve as an indicator of immune activity.

**Figure 2:**
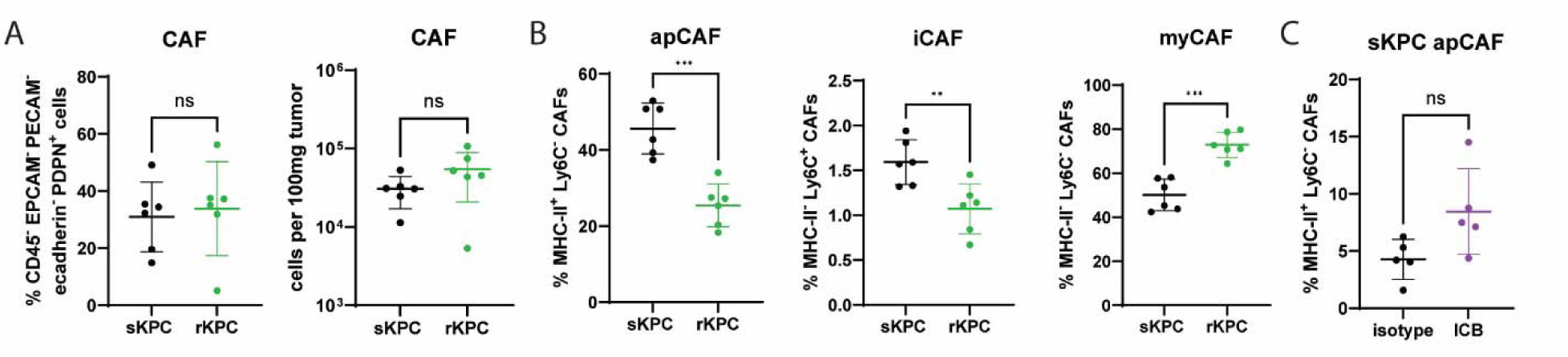
Immunosensitive PDAC TME is characterized by apCAF infiltration. **(A-C)** Total CAFs by percent and absolute count normalized to tumor weight in sKPC and rKPC tumors at day 21. **(A)** Total CAFs defined as CD45^-^ EpCAM^-^ e-cadherin^-^ CD31^-^ PDPN^+^ cells. **(B)** Proportion of CAFs identified as apCAFs (MHC-II^+^ Ly6C^-^), iCAFs (MHC-II^-^ Ly6C^+^), and myCAFs (MHC-II^-^ Ly6C^-^). **(C)** Proportion of total CAFs defined as apCAFs in sKPC tumors treated with ICB or isotype control. Data shown is from one experiment, plotted as mean ± SD; n=5-6 mice per group.

### apCAF-depleted sKPC tumor-bearing mice lose sensitivity to ICB

There are several studies that report opposing observations regarding the functional implications of apCAFs. While initial studies showed that apCAFs induce differentiation of Tregs and inhibit the anti-tumor response, others have demonstrated association of apCAFs with better clinical prognosis and treatment response (8,11–14). The functional role of this cell subset therefore remains unclear. Given the quantitative increase in apCAFs both in T cell high tumors and following ICB, we hypothesized that apCAFs are important in mediating sensitivity of sKPC tumor-bearing mice to ICB. To investigate this further, we utilized two approaches to functionally deplete apCAFs.

First, we generated an apCAF knockout mouse model by utilizing the Cre-lox system to delete MHC-II expression in a pan-fibroblast tagged-Cre mouse. We chose PDGFRα-CRE (C57BL/6-Tg(Pdgfra-cre)1Clc/J), in which Cre recombinase expression is under control of promoter and enhancer elements of the *Pdgfra* gene (20,21), and I-AB-flox (B6.129X1-H2-Ab1tm1Koni/J) mice, containing a *loxP*-flanked neo cassette upstream of exon 1 and a *loxP* site downstream of 1 of exon 1 of the *H2-Ab1* locus (22,23), both purchased from Jackson Labs. Excision of MHC-II in PDGFRα-expressing cells rendered fibroblasts unable to express MHC-II, while preserving other fibroblasts and conventional antigen-presenting cells. We orthotopically implanted sKPC tumors in apCAF I-AB KO and control I-AB-flox mice and treated them with dual ICB for three weeks (**Figure 3A**). There was no difference in CD4 T cells, CD8 T cells, and DCs between apCAF KO and control mice (**Supplementary Figure 2**). Despite an incomplete depletion of MHC-II in apCAFs due to mosaicism in the knockout colony (**Figure 3B**), we observed larger tumors in apCAF KO sKPC-tumor bearing mice as compared with controls (**Figure 3C**), implicating apCAFs as mediators of ICB-sensitivity.

**Figure 3:**
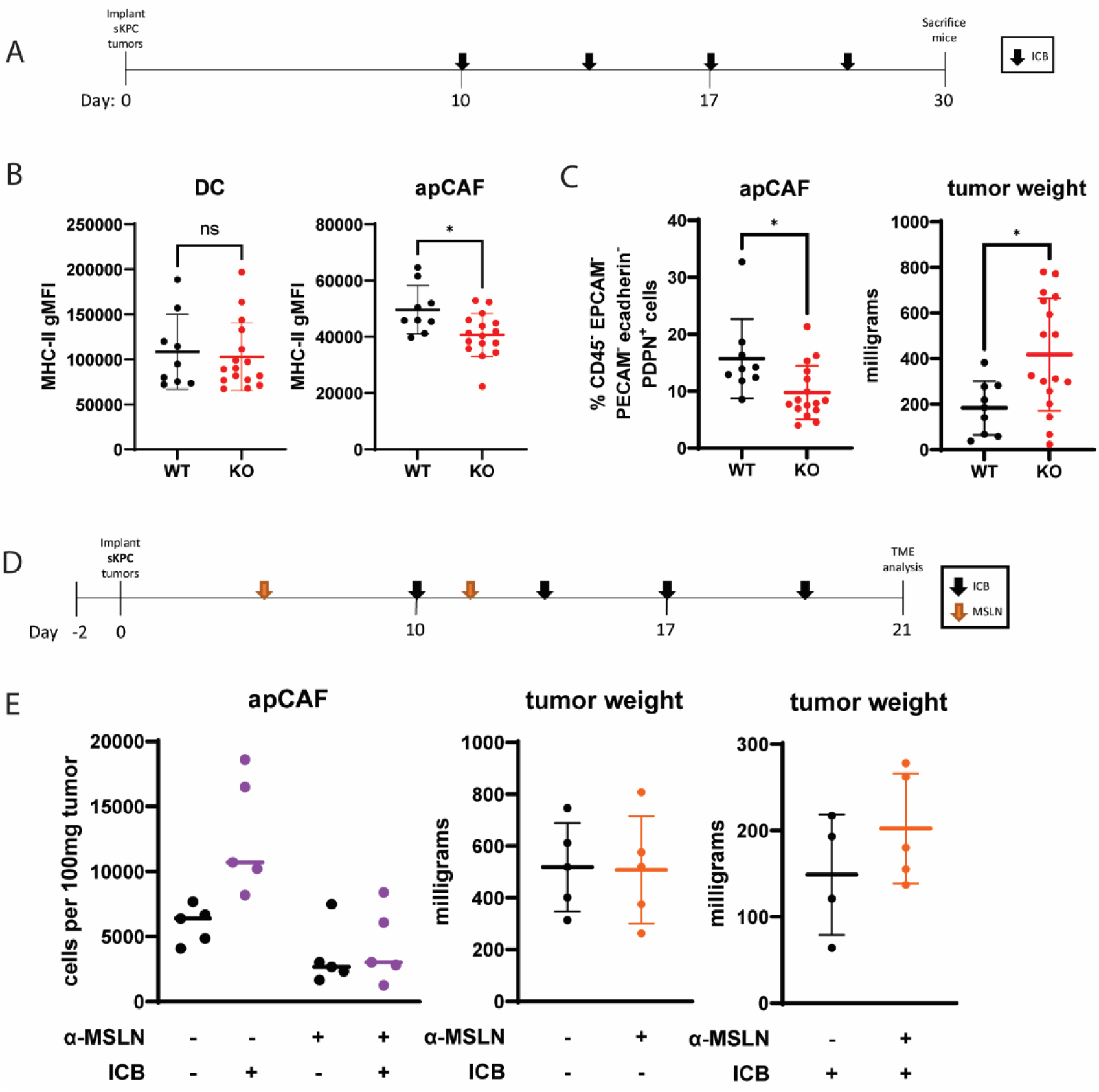
apCAF-depleted sKPC tumor-bearing mice lose sensitivity to ICB. C57BL/6-Tg(Pdgfra-cre)1Clc/J and B6.129X1-H2-Ab1tm1Koni/J breeders were purchased from Jackson Laboratories and bred starting at 10 weeks of age. Litters were genotyped using Transnetyx services. **(A-C)** Experiment timeline **(A)** for ICB treatment of sKPC-bearing apCAF KO mice. Both males and females were used for tumor implantation to maximize the size of available cohorts. **(B)** MHC-II expression on DCs and apCAFs in WT and apCAF KO mice. Data shown are from two independent experiments. **(C)** apCAF quantification and tumor weights in WT and apCAF KO mice. apCAFs are defined as MHC-II^+^ Ly6C^-^ CAFs. **(D-E)** Experiment timeline **(D)** for mesothelin and ICB treatment of sKPC-bearing C57BL/6 mice. **(E)** apCAF quantification (left), and tumor weights for isotype treated (center) and ICI treated (right) C57BL/6 mice depleted of apCAFs with anti-mesothelin antibody. Data shown are from one experiment, n=5 mice per group.

To replicate these results using an alternate method to deplete apCAFs, we used antibody-mediated depletion of the progenitors of apCAFs. Whereas most fibroblasts stem from a mesenchymal cell lineage, apCAFs are derived from the mesothelium (8). We therefore utilized an anti-mesothelin monoclonal antibody to deplete these progenitor cells as previously described (8), and then implanted orthotopic sKPC tumors and treated with dual ICB (**Figure 3D**). Anti-mesothelin treatment yielded an incomplete reduction in apCAFs, yet ICB-treated mice depleted of apCAFs trended towards larger tumors compared with isotype controls (**Figure 3E**). Together, these data demonstrate that despite the fact that apCAFs are only a small population within the PDAC TME, their partial depletion is sufficient to diminish ICB-sensitivity.

### apCAFs colocalize with CD4^+^ T cells

Having established the importance of apCAFs for facilitating sensitivity of sKPC tumors to ICB and hypothesizing that apCAFs mediate this effect via direct interaction with CD4^+^ T cells, we next visualized the localization of these cell subsets within sKPC and rKPC tumors. To visualize apCAFs and CD4^+^ T cells within the TME we utilized imaging mass cytometry (IMC) to analyze the spatial relationship of these cells. Orthotopically implanted sKPC and rKPC tumors were harvested at 10 days post implantation and then formalin fixed and paraffin embedded with the resulting slides stained for IMC. apCAFs and CD4^+^ T cell phenotypes were defined with the markers shown in **Supplementary Tables 1-2**. Phenotypes were identified in all regions of interests, representative images of which are shown (**Figure 4A-B**). We then used the HighPlex FL algorithm in HALO (Indica Labs) and neighboring distance functions to calculate the number of apCAFs within 50 µm of CD4 T cells. This analysis was additionally calculated for apCAFs within 50 µm of Tregs. We observed that apCAFs are enriched within 10-20 µm, approximately one cell diameter, of CD4^+^ T cells (**Figure 4C**) and Tregs (**Figure 4D**) in both sKPC and rKPC tumors, suggesting that these two cell populations interact within the PDAC TME. These analyses also confirmed that there are more apCAFs and CD4^+^ T cells in the TME of sKPC tumors compared with rKPC tumors. Interestingly, we found that apCAFs in sKPC tumors are closer in proximity to CD4^+^ T cells and Tregs, both by absolute numbers and when normalized as proportions of total apCAFs. Importantly, though both apCAFs and CD4^+^ T cells are sparse in the rKPC TME, they demonstrate proximity and thus the potential for interaction and cell-cell communication within the TME.

**Figure 4:**
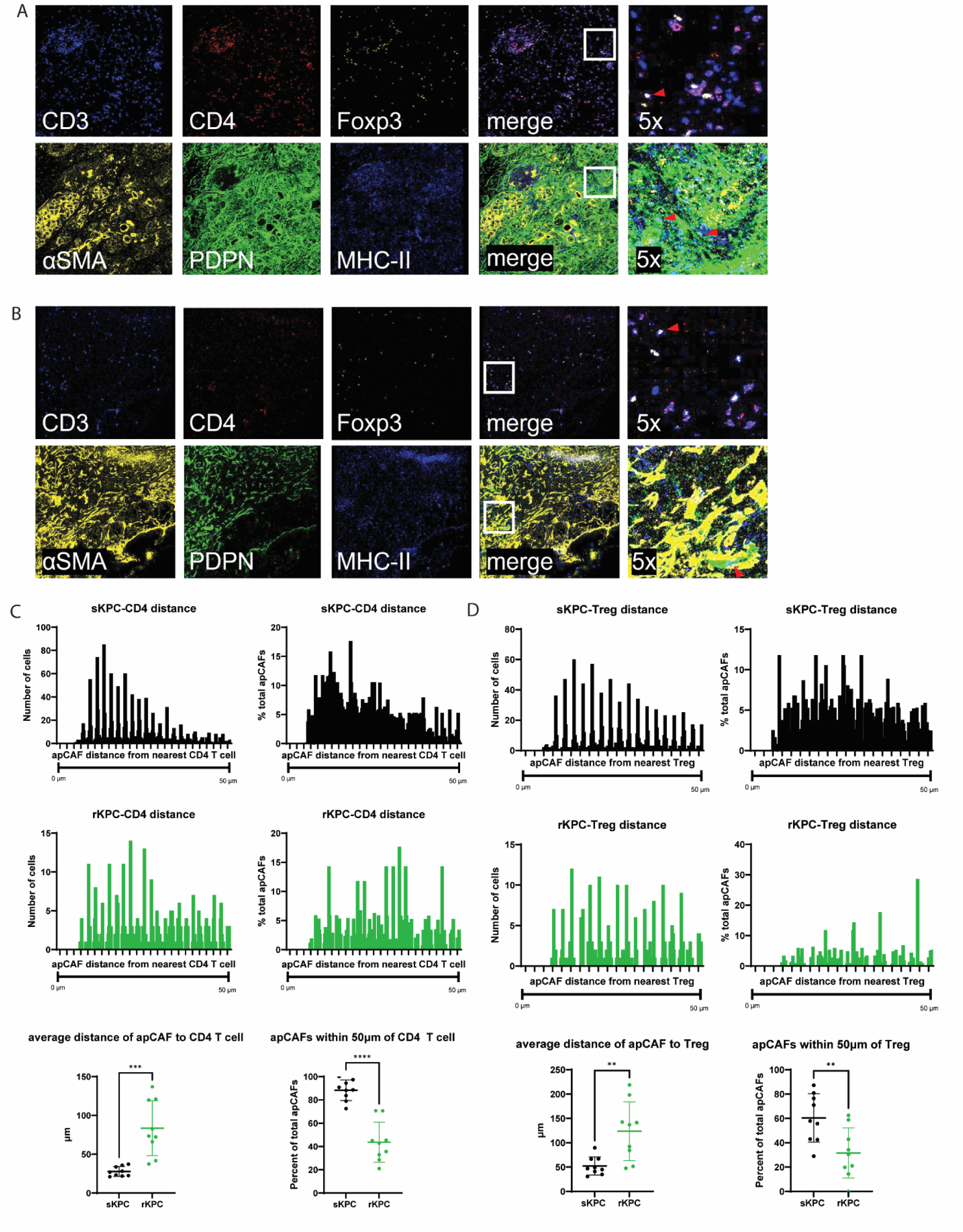
apCAFs are spatially proximal to CD4^+^ T cells. **(A-B)** Representative images of Tregs and apCAFs from sKPC **(A)** and rKPC **(B)** tumors. Tregs were defined by DNA1, CD3, CD4, and Foxp3 expression. apCAFs were defined by DNA1, αSMA, PDPN, and MHC-II expression. Right columns depict the indicated field of view from the merge panel at 5x. Red arrowheads indicate cells matching the phenotype. **(C-D)** Proximity analysis of apCAFs and CD4^+^ T cells **(C)** and Tregs **(D)** was conducted on imaging mass cytometry samples using HALO software. Number (left) and percent of total (right) of apCAFs found in each 2.5µM bin shown.

### apCAFs activate naïve and tumor-infiltrating CD4^+^ T cells

Earlier studies established the functionality of MHC-II on apCAFs by showing that apCAFs pulsed with OVA activate naïve OT-II CD4^+^ T cells by upregulation of cell surface markers (8). To compare apCAF activity between sKPC and rKPC tumors, we similarly tested their influence on naïve OT-II CD4^+^ T cells. apCAFs from sKPC and rKPC tumors were FACS-sorted three weeks post implantation (**Supplementary Figure 3**). Additionally, CD11c^+^ cells were isolated from these same tumors to represent tumor-infiltrating dendritic cell APCs. APCs were pulsed with OVA_323-339_ and cocultured with naïve OT-II CD4^+^ T cells overnight, followed by phenotypic analysis of the T cells (**Figure 5A**). We found that apCAFs from both sKPC and rKPC tumors are equivalent in their ability to induce upregulation of the activation marker CD69 on naïve CD4^+^ T cells, but are not as robust as CD11c^+^ DCs (**Figure 5B-C**). This demonstrates that in an optimal ex-vivo environment, apCAFs function as efficient APCs regardless of tumor of origin.

**Figure 5:**
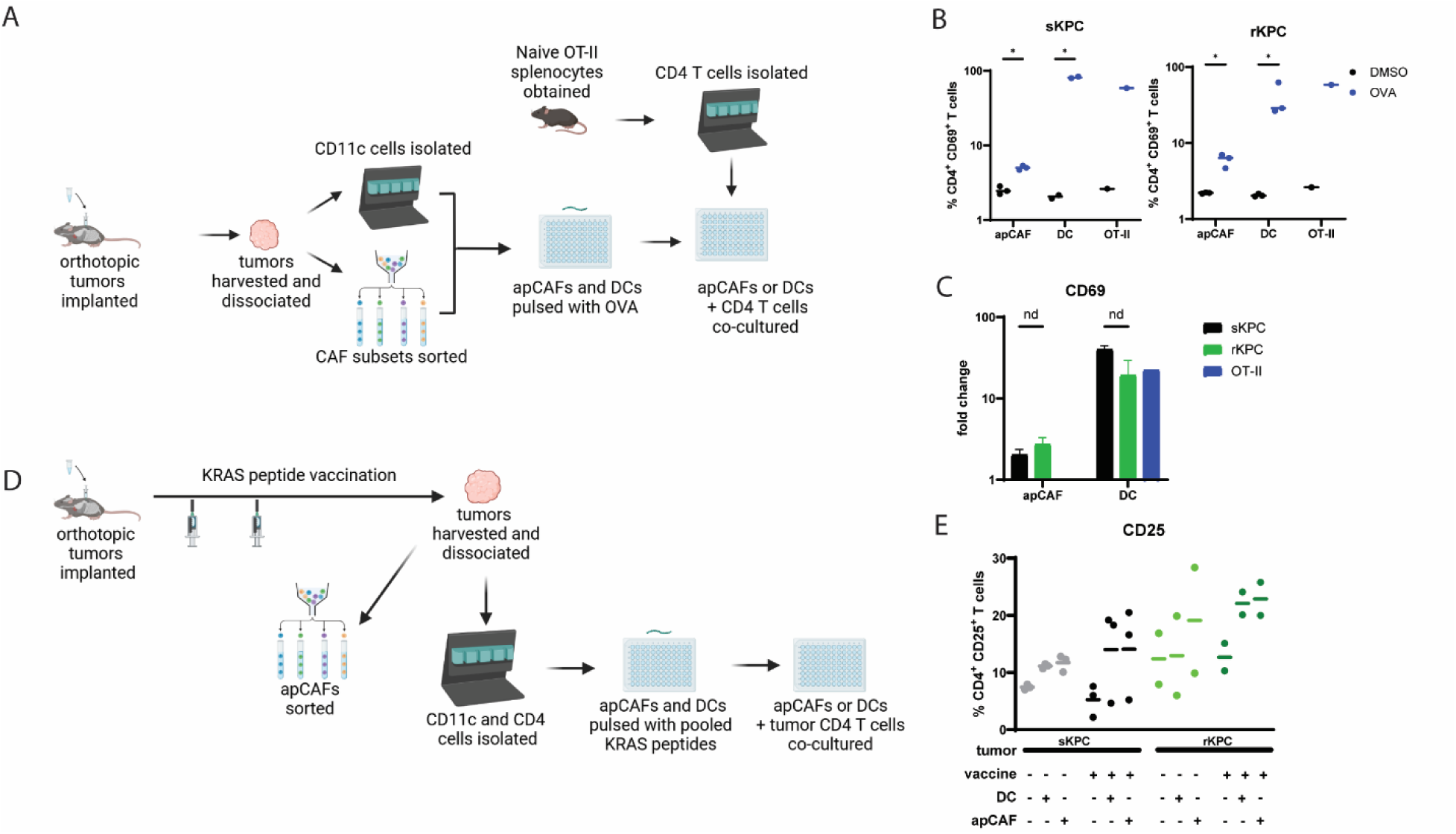
apCAFs activate naïve and tumor-infiltrating CD4^+^ T cells. **(A-C)** Sorted apCAFs and bead-isolated CD11c^+^ cells (DC) from sKPC and rKPC tumors were pulsed with OVA_323-339_ and co-cultured with naïve OT-II CD4^+^ T cells. CD11c^+^ OT-II splenocytes were pulsed with OVA for a positive control (OT-II). **(B)** CD69 expression on CD4 T cells stimulated by apCAFs or DCs from sKPC (left) and rKPC (right) tumors. **(C)** Fold change of CD69 expression comparing CD4 T cells stimulated by apCAFs or DCs pulsed with OVA peptide or with solvent alone. **(D-E)** Sorted apCAFs and bead-isolated CD11c^+^ cells from KRAS-vaccinated sKPC and rKPC tumor-bearing mice were pulsed with KRAS_G12D_ and co-cultured with tumor-infiltrating CD4^+^ T cells. **(E)** CD25 expression on tumor-infiltrating CD4^+^ T cells co-cultured with CD11c^+^ cells or apCAFs under the conditions shown.

Knowing that apCAFs can serve as APCs in an ex-vivo coculture system, we next asked whether apCAFs can also function as APCs for tumor-infiltrating lymphocytes. To investigate this interaction, CD4^+^ T cells were isolated from sKPC tumors in addition to apCAFs and CD11c^+^ DCs. APCs and tumor-infiltrating CD4^+^ T cells were cocultured overnight in the absence of exogenous peptides. Unsurprisingly, analysis of cell surface activation markers CD25 and CD69 revealed no difference regardless of presence of an APC (**Supplementary Figure 4A**). Tumor CD4^+^ T cells already exhibit baseline degree of activation and it is notoriously difficult to elicit antigen-specific responses in the PDAC TME (24,25).

The KPC tumor clones used in this study were generated on a background of KRAS_G12D_ mutations and thus we next pulsed sKPC and rKPC apCAFs and DCs with KRAS_G12D_ peptides prior to overnight coculture with tumor-infiltrating CD4^+^ T cells. Again, these peptide-pulsed APCs did not activate tumor-infiltrating CD4^+^ T cells (**Supplementary Figure 4B**). Given the known paucity of tumor neoantigens in PDAC tumors (26–28), it is unsurprising that there were few KRAS-specific T cells to activate with KRAS_G12D_-pulsed APCs.

One approach to creating a more immunogenic TME is utilization of tumor antigen vaccines (29). Our group has developed a peptide vaccination strategy comprised of the six most common KRAS mutations that prime and boost a KRAS-specific immune response (manuscript in preparation). We thus utilized this vaccination strategy with sKPC and rKPC tumor-bearing mice to elicit an antigen-specific T cell response and cocultured KRAS_G12D_-pulsed APCs with tumor-infiltrating CD4^+^ T cells (**Figure 5D**). With this approach we found that apCAFs were able to elicit upregulation of CD25 activation markers commensurate with what was observed in naïve OT-II CD4^+^ T cells (**Figure 5E**). Together, these results demonstrate that apCAFs not only function as effective APCs for naïve transgenic T cells but also for tumor-infiltrating CD4^+^ T cells, further showing that apCAFs may play a role in facilitating T cell activity within the PDAC TME.

### apCAFs from rKPC tumors induce differentiation of Tregs with greater suppressive signatures

Knowing that apCAFs activate tumor-infiltrating CD4^+^ T cells, we next sought to identify whether this activation led to variability in CD4^+^ T cell differentiation. Prior studies have shown that apCAFs induce differentiation of Tregs from naïve CD4^+^ T cells (8). Following overnight coculture with OVA_323-339_-pulsed apCAFs from rKPC tumors (**Figure 5A**), we observed that 10-15% of naïve OT-II CD4^+^ T cells differentiated into Foxp3^+^ Tregs; interestingly we did not see an increase in Treg differentiation when CD4^+^ T cells were cocultured with sKPC apCAFs (**Figure 6A**). Tumor-infiltrating CD4^+^ T cells from KRAS-vaccinated mice cocultured with KRAS peptide-pulsed apCAFs similarly displayed differentiation to a Treg phenotype irrespective of tumor origin of the apCAF (**Figure 6B**). Given our finding that apCAF-depleted tumor-bearing mice lose partial sensitivity to immune checkpoint blockade, we next asked whether these Tregs were functionally immunosuppressive. Interestingly, we found that following interaction with apCAFs from an sKPC tumor, fewer Tregs produced TGFβ as shown by intracellular cytokine staining and ELISA (**Figure 6C**). Together these data show that although there is an increase in the Treg population following apCAF stimulation, Tregs in sKPC tumors may not be functioning in a classical anti-inflammatory manner.

**Figure 6:**
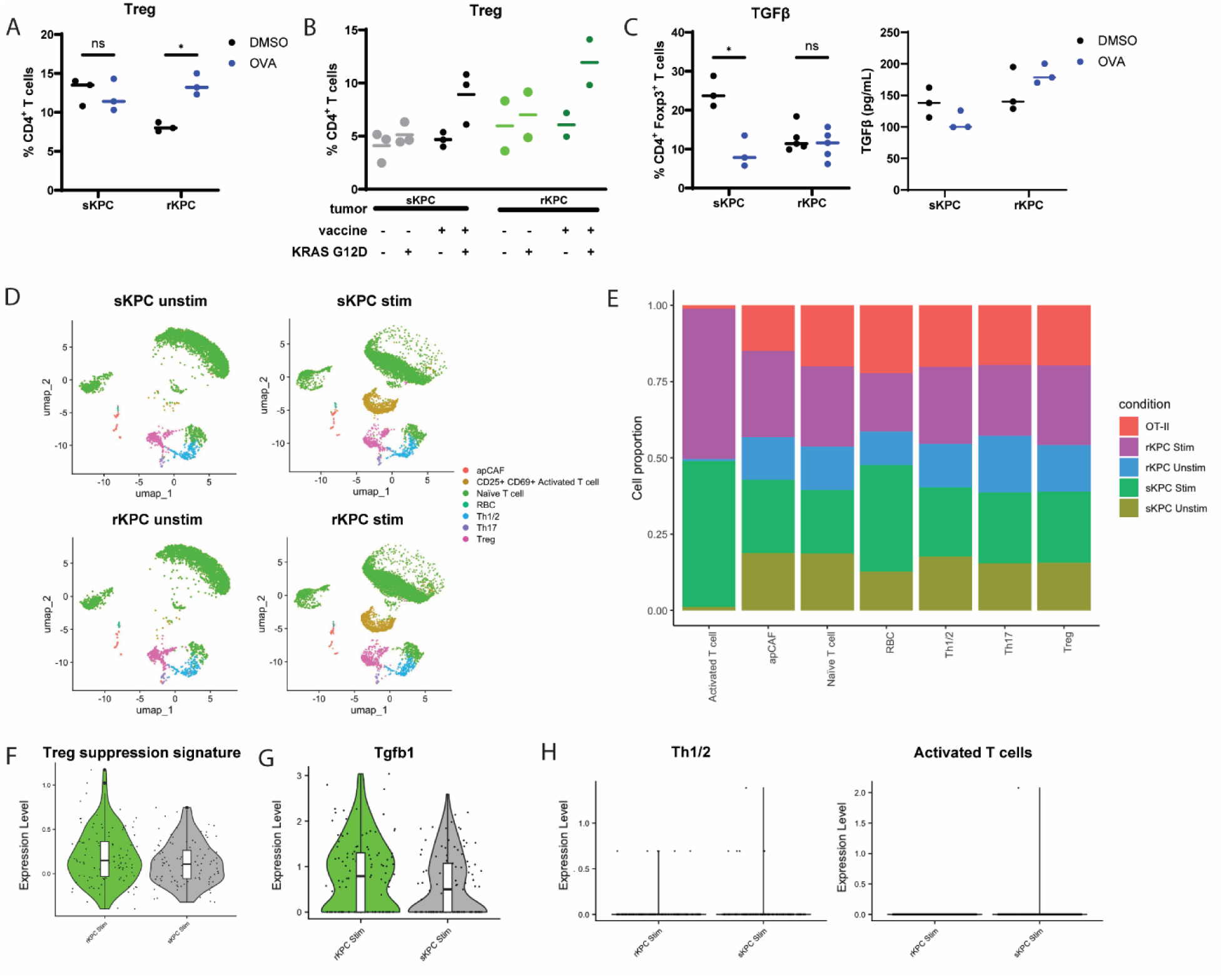
apCAFs from rKPC tumors induce differentiation of Tregs with greater suppressive signatures. **(A)** Sorted apCAFs from sKPC and rKPC tumors were pulsed with OVA_323-339_ and co-cultured with naïve OT-II CD4^+^ T cells; quantification of Foxp3^+^ Tregs. **(B)** Sorted apCAFs from sKPC and rKPC tumors following vaccination were pulsed with KRAS peptides and co-cultured with tumor-infiltrating CD4^+^ T cells; quantification of Foxp3^+^ Tregs. **(C)** Intracellular cytokine staining (left) and ELISA (right) for TGFβ following co-culture with apCAFs. **(D-H)** apCAFs pulsed with OVA_323-339_ were co-cultured overnight with naïve OT-II cells and processed for single-cell RNA sequencing. **(D)** umaps of apCAF and CD4^+^ T cell clusters. **(E)** Proportion of cells representing each CD4^+^ T cell or apCAF cluster in the indicated experimental conditions. **(F)** Expression of a Treg suppression signature consisting of *Il2ra*, *Il10*, *Ctla4*, *Ebi3*, *Tgfb1*, *Ccr4*, *Tbx21* in Tregs. **(G)** Expression of *Tgfb1* in Tregs. **(H)** Expression of *Ifng* in Th1/2 and activated T cell clusters.

To understand whether the difference in TGFβ production from Tregs stimulated by sKPC and rKPC apCAFs reflected changes in Treg phenotype and canonical Treg cell suppression mechanisms, we again cultured apCAFs from sKPC and rKPC tumors with naïve OT-II CD4^+^ T cells in the presence or absence of OVA_323-339_ and performed single-cell RNA-sequencing. All CD4^+^ helper T cell states were represented in the data (**Figure 6D**). Tregs were represented in all conditions, with expansion of the population in the peptide-stimulated conditions (**Figure 6E**).

Treg suppression mechanisms vary depending on the environmental context and are not dependent on just one set of cytokines, such as TGFβ or IL-10, but also include *Ctla4*, *Il2ra*, *Ebi3*, and *Ccr4* among others (30–33). We therefore combined these markers into a “Treg suppression” gene set and observed higher expression of this suppressive signature in Tregs stimulated by rKPC apCAFs (**Figure 6F, Supplementary Figure 5**). Furthermore, in line with differential TGFβ production that we observed in our flow cytometry experiments, we found higher TGFβ gene expression in Tregs cocultured with apCAFs from rKPC tumors (**Figure 6G**). Conversely, analysis of the Th1/2 and activated T cell clusters showed higher *Ifng* expression in T cells stimulated by sKPC apCAFs (**Figure 6H**). Together, these data suggest that rKPC apCAFs induce Tregs with higher expression of cell suppression mechanisms that dampen the effector immune response.

### apCAF – CD4^+^ T cell signaling varies with tumor origin of apCAF

We next performed differential gene expression analysis on CD4^+^ T cell clusters to investigate further transcriptional differences based on tumor of origin. We found very few genes or pathways differentially expressed within all CD4^+^ T cells or within Treg clusters between the sKPC or rKPC conditions, nor when comparing Tregs in apCAF-pulsed and -unpulsed conditions. (**Figure 7A-D, Supplementary Figure 6**). However, the difference in Treg phenotype and function induced by apCAFs from the sKPC and rKPC TME suggests these apCAFs exhibit differential signaling to CD4^+^ T cells influenced by the different environment from which they were derived. To look more closely at signaling between apCAFs and CD4^+^ T cells, we performed cell to cell communication inference using dominoSignal (34,35) from apCAFs to each CD4^+^ T cell cluster. This network-based communication inference algorithm identifies communication based on the association of receptor expression with transcription factor activity, quantified using SCENIC (36), as a measure of receptor activation in receiver cells as well as ligand expression by sender cells. dominoSignal identifies signaling on the basis that (1) the receptor is reliably expressed in the receiver, (2) the ligand is expressed in the sender, and (3) the transcription factors linked to receptors are differentially active in the receiver cell type. Strikingly, certain signaling pathways in CD4^+^ T cells were differentially represented depending on apCAF tumor of origin (**Supplementary Figure 7**). When focusing on apCAF signals towards Tregs, we observed a unique pattern of signaling involving *Cxcl9*, *Ccl22*, *Il6*, and *Cd274* (PD-L1). We found that rKPC apCAFs signaled to all CD4^+^ T cell clusters via *Cd274* (PD-L1) to *Pdcd1* (PD-1), *Il6* to *Il6ra*, and *Ccl22* to *Cd26* (Dpp4) axes (**Figure 7E-F**), all of which are associated with dampening of the effector immune response and with aggressive tumor growth (37–42). Conversely, sKPC apCAFs signaled to CD4^+^ T cell clusters via *Cxcl9* to *Cxcr3* and *Dpp4* receptors (**Figure 7E-F**); CXCL9-CXCR3 interactions have been shown to be enriched in ICB responders (43–45). In the Tregs and Th1/2 clusters, both CXCR3 and Dpp4 receptors were predicted to be engaged by CXCL9, implicating competition for Dpp4 receptors on receiver cells by both chemokines.

**Figure 7:**
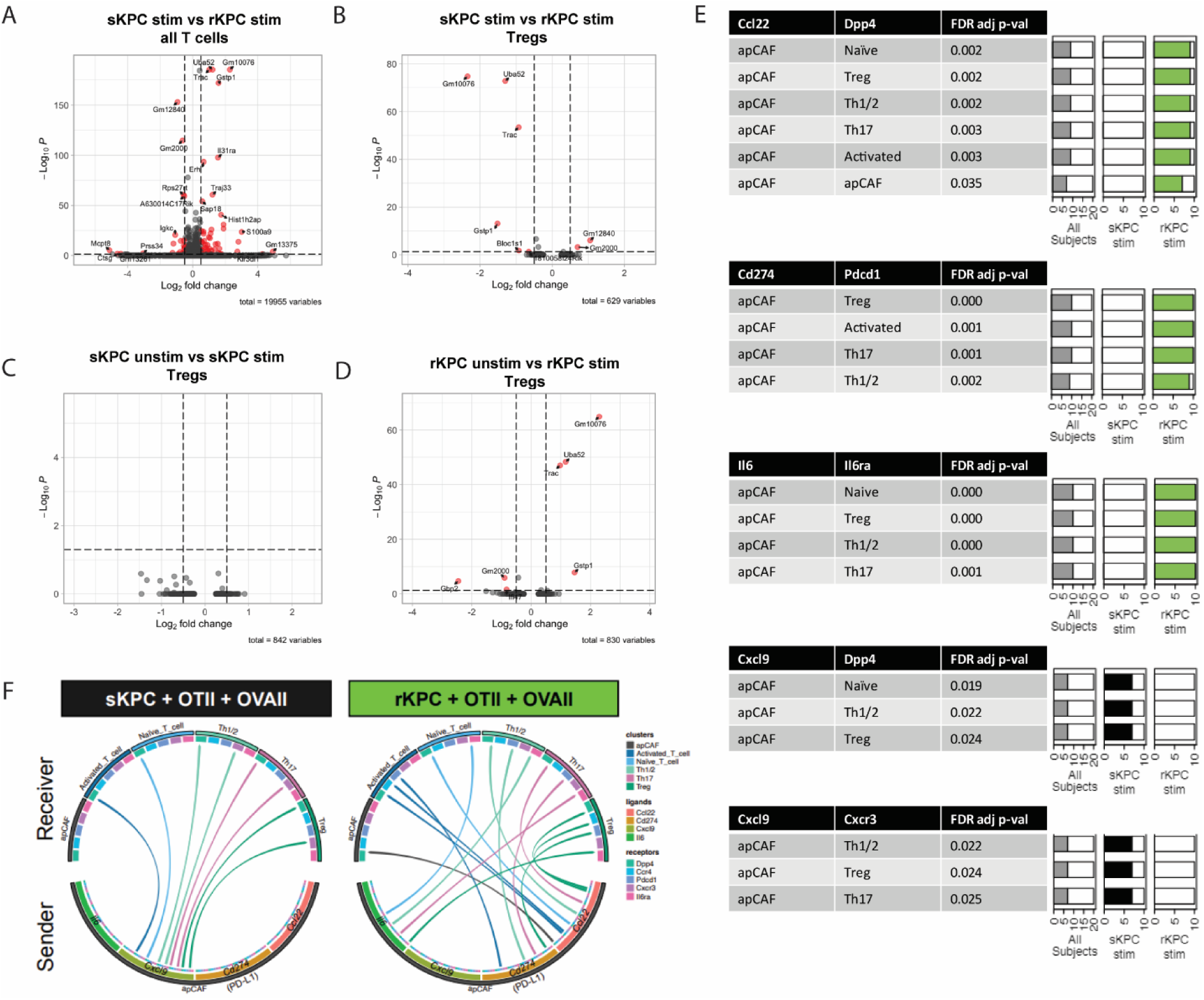
apCAF – CD4 signaling varies with tumor origin of apCAF. **(A-D)** Volcano plots showing differentially expressed genes in CD4^+^ T cells cocultured with apCAFs from sKPC or rKPC tumors. **(A)** DEG of all CD4^+^ T cells cocultured with pulsed sKPC apCAFs (right) versus pulsed rKPC apCAFs (left). **(B)** DEG of Tregs cocultured with pulsed sKPC apCAFs (right) versus pulsed rKPC apCAFs (left). **(C)** DEG of Tregs cocultured with unpulsed (right) versus pulsed (left) sKPC apCAFs. **(D)** DEG of Tregs cocultured with unpulsed (right) versus pulsed (left) rKPC apCAFs. **(E-F)** 10 bootstrapped samples of apCAFs and CD4^+^ T cells were generated from sKPC and rKPC experiments. Each bootstrapped sample was subjected to cell-cell communication inference using dominoSignal (v 1.1.0) and the dependence of inferred ligand-receptor signals from apCAFs to functional CD4 T cell subsets upon whether cells came from sKPC or rKPC tumors was tested by Fisher’s Exact test. The leading differential signals using Il6, Cxcl9, Cd274 (PD-L1), and Ccl22 are shown. **(E)** Table depicts sender cell and ligand (left), receiver cell and receptor (center), and FDR adjusted p-value of the Fisher’s exact test for occurrence of intercellular signals being inferred as active in each tumor type (right). The bars on the right show the proportion of all bootstrapped samples (grey), sKPC samples (black), and rKPC samples (green) where the signal was inferred as active by dominoSignal. **(F)** Circos plot depicting selected signaling pathways. The lower semicircle represents ligands sent by apCAFs and the upper semicircle represents receptors on recipient cell types. Outer arcs represent cell types and inner arcs represent ligands and receptors. Chords are draws between ligands and receptor signals that differentially occurred in one tumor type and are colored by recipient cell type.

## Discussion

Therapeutic strategies to target CAFs and influence anti-tumor activity have focused on CAFs as a homogeneous population and shown little clinical benefit (5,46–51). With a developing understanding of heterogeneity in CAF phenotype and function, it is important to understand the differing roles of CAF subsets. To date, apCAFs have not been evaluated as targets for immune modulation. Given their distinct origin and their unique ability to interact with T cells, further understanding of their role in influencing immune responses should lead to more specific apCAF modalities for treatment.

This study makes several contributions to our understanding of apCAF biology and reveals potential avenues for translation. First, we have shown a quantitative difference in apCAFs that relates to T cell activity in the TME and tumor burden, suggesting that increasing apCAF prevalence within the TME could improve anti-tumor activity. Further characterization of apCAFs in human PDAC tissue may reveal apCAFs as predictive biomarkers or indicators of response, as has been previously suggested (13,15). Second, we show that regardless of TME of origin, apCAFs can similarly interact with and stimulate CD4^+^ T cells in an ex-vivo experimental setup, suggesting that even apCAFs from an immunoresistant TME are capable of promoting effector cell responses if given the right environmental cues. While modulating CAFs as one homogeneous population has not yielded clinical benefit, specific modulation of apCAFs may promote an effector immune response; it will be important to study differences in apCAFs between varying TMEs to better understand apCAF differentiation. Finally, we see that apCAFs have a qualitative difference in the ability to induce Treg differentiation both phenotypically and functionally via differential chemokine signaling dependent on the TME of origin, which reveals specific ligand-receptor interactions to modulate and increase anti-tumor activity in a resistant tumor. This also emphasizes the need to further interrogate the interactions between apCAFs and other CD4^+^ T cell subsets as well as with other immune cells in the TME.

IMC data confirmed apCAFs are in close proximity with CD4^+^ T cell subsets within tumors and thus have the potential to interact within the TME. This has previously been shown in precancerous lesions (10) and suggested in tumor biopsies from melanoma, colorectal cancer, and gastric cancer patients (15,52). Coculture experiments confirmed apCAFs can present antigen to both transgenic and tumor-infiltrating CD4^+^ T cells in an antigen-specific manner and induce Treg differentiation. However, variability in Treg phenotype and function was dependent on the TME from which apCAFs were derived, and showed diminished TGFβ production and increased T cell activating chemokine signaling when derived from ICB sensitive tumors. In the context of an isolated interaction with apCAFs from the rKPC TME, we show here that naïve CD4^+^ T cells adopt a stronger suppressive phenotype which also correlates with weaker effector cytokine expression in Th1 cells.

Our findings of differential chemokine signaling from apCAFs towards Tregs reinforces and emphasizes that each unique TME can have a different impact on cellular interactions, even in the absence of differential expression of individual receptors or ligands. apCAFs from sKPC tumors had higher expression of CXCL9 signaling towards Tregs. CXCL9, also known as monokine induced by gamma interferon (MIG), signals via the CXCR3 receptor and is a known chemoattractant (37,53), which may be an explanation for our observation that apCAFs and CD4^+^ T cells were in closer proximity in sKPC tumors. Furthermore, CXCR3-CXCL9 interactions are enriched in patients who are ICB responders (43–45). Conversely, apCAFs from rKPC tumors had higher expression of CCL22 signaling towards Tregs via Dpp4 receptors, as well as higher levels of IL6 and PD-L1 signaling as compared with apCAFs from sKPC tumors. CCL22 signaling is strongly associated with Treg suppressive activity and has been shown to promote tumor growth and escape in several different models (37–42). Dpp4, which can be either soluble or a membrane-bound receptor, is closely related to fibroblast activation protein (FAP) which has a strong association with worse prognosis in PDAC (54,55). Dpp4 inhibition has been shown to enhance both NK and T cell immune infiltration in murine models of PDAC via recruitment through increased CXCL9 expression (56). Similarly, IL-6 signaling is thought to accelerate tumor progression by promoting differentiation and expansion of immunosuppressive cells (57–59).

It is important to note that the single-cell RNA sequencing experiments in this study were performed with naïve OT-II CD4^+^ T cells rather than tumor-infiltrating CD4^+^ T cells or cells from an otherwise inflammatory environment. This eliminates existing heterogeneity of Tregs within the TME, which could represent peripheral (pTreg) or thymic (tTreg) Tregs that could differ between the sensitive and resistant TME (60). While this helps us eliminate variability in existing Treg differentiation influenced by the TME and isolates the effect of apCAFs from sKPC and rKPC tumors on naïve CD4^+^ T cells, we recognize that this may not reflect the extent of intratumoral apCAF-Treg interactions. The PDAC TME is highly multicellular outside of just CD4^+^ T helper cell states; in this study we have isolated one set of cellular interactions for in-depth studies but cannot account for the influence on these interactions by other stromal, immune, and tumor cells.

There are many technical challenges when attempting to deplete rare cell populations, such as apCAFs, in vivo. Both the Cre-Lox system and intraperitoneal administration of monoclonal antibodies targeting an apCAF marker, mesothelin, resulted in incomplete functional depletion of the apCAF population. However, even without robust depletion of apCAF function, we observed a relative increase in tumor weight in sKPC tumor-bearing mice subjected to both methods of depletion. Despite the limitations of both techniques, the consistent diminished response to ICB in apCAF-depleted sKPC tumor-bearing mice strongly suggests that apCAFs are positively associated with an anti-tumor response. Additionally, the Cre-Lox approach does not eliminate apCAFs as a population but rather prevents these cells from MHC-II -mediated mechanisms of interaction with target cells. We cannot discount the possibility that apCAFs may also be functioning through MHC-II -independent pathways to exert influence.

In summary, our findings demonstrate a role for apCAFs in modulating the anti-tumor response in PDAC tumors. Studies featuring congenic murine PDAC tumor clones have clearly demonstrated that tumor-cell intrinsic factors shape the TME and influence response to immunotherapy; here we see that chemokine signaling from apCAFs similarly shapes the TME and influences therapeutic response. Understanding the scope of function of this unique stromal cell subset and their regulation will facilitate the development of more effective anti-tumor immune based approaches for PDAC patients.

## Materials & Methods

### Cell lines

2838c3 and 5419c5 KPCY cell lines were generously provided by Dr. Benjamin Stanger’s laboratory (University of Pennsylvania, Philadelphia, PA). Cells were maintained in DMEM (catalog) supplemented with 10% heat-inactivated fetal bovine serum (catalog) and 1% penicillin and streptomycin (catalog). Cells were grown at 37°C with 10% CO2 in a humidified incubator.

### Animal experimentation

All mice used in this study were male C57Bl/6J mice purchased from Jackson Laboratories. Mice were 8 weeks of age at time of the initiation of experiments. Female mice were utilized for all experiments unless otherwise specified. Mice were maintained in accordance with the Institutional Animal Care and Use Committee (IACUC) guidelines. Mice were fed a standard diet, were not fasted prior to the initiation of an experiment or an assessment, and all interventions were performed during the light cycle. C57BL/6J (JAX #000664), B6.Cg-Tg(TcraTcrb)425Cbn/J (JAX #004194), C57BL/6-Tg(Pdgfra-cre)1Clc/J (JAX #013148), and B6.129X1-H2-Ab1tm1Koni/J (JAX #013181) were purchased from The Jackson Laboratory.

#### Orthotopic pancreas injections

The mouse pancreatic orthotopic injection model was performed as described previously(19). Briefly, mice were anesthetized, the abdomen was opened, and 1×10^5^ tumor cells suspended in 50µl of a 95% Geltrex LDEV-Free Reduced Growth Factor Basement Membrane Matrix (Thermo Fisher, catalog # A1413201) and 5% PBS solution were injected into the pancreas.

#### Survival analyses

Mice harboring orthotopic tumors were monitored throughout experiments. Mice were euthanized if they were exhibiting signs of poor condition including lethargy, hunched posture, or significant abdominal distension per institutional guidelines. All euthanized mice were examined to confirm presence of pancreatic tumors. Kaplan-Meyer analyses of survival were used to visual the survival curves with significant differences between groups determined by log-rank testing.

#### In vivo treatments

For immune checkpoint blockade experiments, intraperitoneal injections of 200 µg anti-CTLA4 (clone 9H10, BioXCell, catalog # BE0131) or polyclonal Syrian hamster IgG (BioXCell, catalog # BE0087) once on day 10 and 200 µg anti-PD1 (clone RMP1-14, BioXCell, catalog # BE0146) or rat IgG (clone 2a3, BioXCell, catalog # BE0089) antibodies twice weekly were administered starting on day 10. For apCAF depletion experiments, 50 µg anti-mesothelin antibody (MBL Life Science) was administered intraperitoneally prior to surgery and weekly thereafter.

KRAS peptide vaccines were prepared immediately prior to injections. Briefly, STING agonist ADU-S100 (InvivoGen, catalog # tlrl-nacda2r-1)was diluted to 1 µg/µl in PBS and warmed to 37C for 15 minutes, followed by addition of 50µg of each KRAS peptide (G12D, G12C, G12R, G12A, G12V, and G13D). Mice were vaccinated with KRAS vaccine or PBS twice, 7 days apart. Vaccine was delivered in 100µl volume subcutaneously at the base of the tail (*manuscript and rationale for this protocol in preparation*).

#### apCAF KO generation

C57BL/6-Tg(Pdgfra-cre)1Clc/J and B6.129X1-H2-Ab1tm1Koni/J were crossed to generate a Pdgfra^Cre^;H2Ab1^flox/flox^ apCAF KO strain. Genotyping of all mice was conducted through Transnetyx, Inc.

#### Mouse tumor dissociation

Tumors were dissociated to a single-cell suspension using a mouse tumor dissociation kit (Miltenyi Biotec, catalog # 130-096-730) with the gentleMACS dissociator system. Tumors were mechanically chopped into small fragments and transferred to a Miltyeni gentleMACS C tube (Miltenyi Biotec, catalog # 130-093-237) along with dissociation enzymes. Tumors were digested in the gentleMACS dissociator with the mTDK2 setting. Once dissociated, tumors were filtered twice using a 100 μm filter (CELLTREAT, catalog # 229485) followed by a 70 μm filter (CELLTREAT, catalog # 229483). Single-cell suspensions were centrifuged at 1500 rpm for 5 minutes; supernatant was aspirated and then resuspended in 1 mL ACK lysis buffer for 3 minutes at room temperature, then quenched with 10ml FBS-containing media. Cells were centrifuged at 1500 rpm for 5 minutes; supernatant was aspirated and resuspended in DMEM with 10% FBS.

#### Cell isolations

apCAFs were FACS-sorted using the BD Fusion sorter. Single-cell tumor suspensions were pooled from up to five tumors and stained with Zombie NIR live/dead stain, CD45, CD31, e-cadherin, EpCAM, PDPN, MHC-II, and Ly6C. apCAFs were defined as CD45^-^ CD31^-^ e-cadherin^-^ EpCAM^-^ PDPN^+^ MHC-II^+^ Ly6C^-^ cells. Cells were collected into cold PBS with 2% FBS.

CD11c^+^ and CD4^+^ T cells were isolated from single-cell tumor suspensions using CD11c MicroBeads UltraPure (Miltenyi Biotec, catalog # 130-125-835) and CD4 (TIL) MicroBeads (Miltenyi Biotec, catalog # 130-116-475) according to manufacturer’s protocol.

Splenocytes from naïve OT-II mice were processed by mashing spleens through a 40 μm filter (CELLTREAT, catalog # 229481). Single-cell suspensions were centrifuged at 1500 rpm for 5 minutes; supernatant was aspirated and then resuspended in 1 mL ACK lysis buffer for 3 minutes at room temperature, then quenched with 10ml FBS-containing media. Cells were centrifuged at 1500 rpm for 5 minutes; supernatant was aspirated and resuspended in DMEM with 10% FBS. CD4^+^ T cells were then isolated using the EasySep Mouse CD4^+^ T cell Isolation Kit (STEMCELL Technologies, catalog # 19852) per manufacturer’s protocol.

### Flow cytometry

#### APC – T cell coculture

2×10^4^ apCAFs or CD11c^+^ cells were plated and pulsed with 25 µg/ml of OVA_323-339_ or DMSO vehicle control for four hours. Peptides were washed out and 50,000 CD4^+^ T cells were added in fresh medium and cultured overnight. For intracellular cytokine staining experiments, protein transport inhibitor (eBioscience, catalog # 00-4980-03) was added for the last four hours of incubation.

#### Flow Staining

Single-cell suspensions were plated at a density of 1×10^5^ per well for staining unless otherwise noted. Cells were washed twice with PBS and stained for viability in 100 μL of Zombie NIR fixable live/dead stain (1:1500, BioLegend, catalog # 423106) and Mouse BD Fc Block (1:50, BD Biosciences, catalog # 553141) in PBS for 20 minutes at room temperature. Cells were washed twice with FACS buffer (1× HBSS, 2% FBS, 0.1% Sodium Azide), stained with 100 μL extracellular antibody mix diluted in FACS buffer for 20 minutes at 4°C, and again washed twice with FACS buffer. For intracellular cytokine detection, cells were permeabilized in 100 μL fix/perm (Invitrogen, catalog # 00-5523-00) for 10 minutes at room temperature, washed twice in 1× perm/wash buffer, and then stained with 100 μL intracellular antibody mix diluted in 1× perm/wash buffer for 30 minutes at 4°C. Cells were then washed twice with 1× perm/wash buffer and run on a Cytek Aurora flow cytometer. All flow cytometry data were analyzed using FlowJo version 10.10.0.

#### Antibodies

The following antibodies were used for flow cytometry analysis: Zombie NIR Fixable Viability Kit (BioLegend, catalog #423105); CD45-BV510 (clone 30-F11, 1:200, BioLegend, catalog #103138); CD45-APC Cy7 (clone 30-F11, 1:200, BioLegend, catalog #103116); e-cadherin-BV421 (clone DECMA-1, 1:200, BioLegend, catalog #147319); EpCAM-BV421 (clone G8.8, 1:200, BioLegend, catalog #118225); CD31-BV421 (clone 390, 1:200, BioLegend, catalog #102424); PDPN-PECy7 (clone 8.1.1, 1:400, BioLegend, catalog #127412); I-A/I-E (MHC-II)-BV785 (clone M5/114.15.2, 1:200, BioLegend, catalog #107645); Ly6C-BV605 (clone HK1.4, 1:200, BioLegend, catalog #128036); CD4-BV510 (clone GK1.5, 1:200, BioLegend, catalog #100449); CD8-BUV805 (clone 53-6.7, 1:200, BD Biosciences, catalog #612898); CD25-BUV395 (clone PC61, 1:200, BD Biosciences, catalog #564022); CD69-BV711 (clone H1.2F3, 1:200, BioLegend, catalog #104537); Tbet-PECy7 (clone 4B10, 1:100, BioLegend, catalog #644824); Foxp3-PE-CF594 (clone R16-715, 1:100, BD Biosciences, catalog #567373); LAP(TGFβ1)-APC (clone TW7-16B4, 1:100, BioLegend, catalog #141406); F4/80-BV650 (clone BM8, 1:200, BioLegend, catalog #123149); Ly-6G-Pacific Blue (clone 1A8, 1:200, BioLegend, catalog #127612); CD11c-BUV737 (clone HL3, 1:200, BD Biosciences, catalog #612796).

### Imaging Mass Cytometry

#### Staining

Murine pancreatic tumors were formalin fixed and paraffin embedded then cut onto slides. Slides were dewaxed in xylene for 20 minutes then rehydrated in a descending alcohol gradient. Slides then underwent antigen retrieval using Cell Conditioning Solution, CC1 (Roche; catalog # 5279801001) for 1 hour at 105°C followed by blocking in 3% BSA in Maxpar PBS (Standard BioTools; catalog # 201058) for 45 minutes at room temperature in a hydration chamber. Next, slides were stained overnight in a hydration chamber at 4°C with the antibody cocktail listed in Supplementary Table 1. Slides were then stained with Cell-ID Intercalator-Ir (Standard BioTools; catalog # 201192B) for 30 minutes at room temperature in a hydration chamber to stain DNA.

#### Data acquisition and analysis

Slides were ablated using a Hyperion Imaging System (Standard BioTools). Resulting MCD files were converted into ome.tiff format using MCD Viewer software (Standard BioTools; version 1.0.560.6). These files were then analyzed using the HighPlex FL (version 4.2.14) algorithm in HALO (Indica Labs; version 3.6.4134.396). Briefly, cells of interest were defined by the following markers: CD3, CD4 (Th); CD3, CD4, Foxp3 (Treg); aSMA, PDPN, MHCIII (apCAF) with Iridium (“DNA1”) used as a nuclear marker to identify all cells (Supplementary Table 2). Algorithm parameters were manually adjusted for each marker to optimally detect signal with minimal non-specific signal detection. The HighPlex FL algorithm allows for cell data to be stored as objects with X and Y coordinates retained to enable spatial analysis. From the object data, spatial dot plots of the cells of interest were created followed by proximity analysis to quantify distances between cells within 50 µm. Distance data were plotted in histograms with 2.5 µm bins.

### Single-cell RNA sequencing

#### Library prep

The single-cell suspensions from the in vitro co-culture and implanted tumors were the input for the single-cell RNA-sequencing. The initial cell input from each was XXX cells. Single-cell RNA-sequencing library preparations were performed using the Chromium Single Cell 3′ Library and Gel Bead Kit (10x Genomics) followed by sequencing on the NovaSeq 6000 (Illumina), following manufacturer’s recommendations.

#### Data processing

The raw single cell transcriptomic sequencing data was converted from FASTQ files to genes-by-cell count matrices using Cell Ranger (v7.1.0) (10x Genomics). All subsequent data processing was conducted in R version 4.3.2 (61). The Seurat (v5.1.0) R/Bioconductor package was used for the analysis. During data quality check and filtering, only cells with at least 200 genes expressed and no more than 6000, less than 25% mitochondrial reads were used for the downstream analysis. To account for differences in sequencing depth and technical artifact, normalization, variable feature identification and scaling of the filtered count data was done via the gamma–Poisson generalized linear model SCTransform (v0.4.1). Dimensionality reduction was performed via principal component analysis (PCA) on the normalized and scaled counts followed by unsupervised clustering with the Louvain algorithm on the PCA embedding. Clusters were visualized in the two-dimensional space using Uniform Manifold Approximation and Projection (UMAP). Manual curation and annotation of cell types was performed using a combination of curated gene signature profiling through the AddModuleScore function, differential expression testing using the MAST test. Differentially expressed genes were visualized using the R package EnahncedVolcano (v1.20.0) (62).

#### Assessment of differential signaling

Assessment of differential signaling to CD4 T cells between sKPC apCAFs and rKPC apCAFs was conducted in R (v4.2.0). Bootstrap replicates were generated from cells in the sKPC stimulated (SM5H1) and rKPC stimulated (SM7H3) conditions by drawing cells with replacement until as many cells from each cell type in the parent sample were obtained and subjected to cell-cell communication inference (CCI) using dominoSignal. Transcription factor activity scores were quantified using pySCENIC (v0.11.0) with cisTarget mm10 reference transcription factor binding motifs. Mouse ligand-receptor pairs were obtained from CellTalkDB. Bootstrapping of inferred signaling was accomplished by sampling cells from each cell type cluster with replacement until as many cells as that were in the parent sample were obtained. For each condition, 15 bootstraps were generated. The bootstrapped cells underwent CCI using dominoSignal (v1.0.0) with parameters setting receptor-transcription factor linkage at spearman correlation > 0.15, transcription factor assignment to cluster at p < 0.001, and minimum threshold percentage of recipient cells expressing receptor at 5%. Ligand sender clusters were identified as clusters with mean scaled expression of the ligand > 0, and intercellular linkages of recipient cell receptors and sender cell ligands were compiled based on recipient cell type for each bootstrap. Differential signaling between sKPC and rKPC conditions was assessed by fisher’s exact test for the dependence of intercellular signal occurrence upon whether the bootstrap was derived from sKPC cells or rKPC cells. Fisher’s exact test p-values were adjusted for multiple testing accounting for the number of possible ligand-receptor interactions received by a cell type using Benjamini-Hochberg FDR adjustment as implemented in the R stats package. The leading interactions outgoing from apCAFs via ligands *Ccl22*, *Cd274*, *Cxcl9*, and *Il6*, and were displayed using ComplexHeatmap to show the FDR-adjusted fisher’s exact test p-value alongside the proportion of bootstraps from each group with active signaling. Differential signals via these ligands were displayed as a circos plot using circlize.

R code used to conduct assessment of differential signaling is available at https://github.com/FertigLab/apCAF_CD4T_crosstalk.

### Quantification and Statistical Analysis

Statistical analysis was performed with Prism 10 software using unpaired Students t-test. P values < 0.05 were considered significant. Data are represented as mean ± SD unless otherwise noted. Significance was designated as follows: *P < 0.05; **P < 0.01; ***P < 0.001, ****P < 0.0001.

## Data Availability

The data generated in this study are available upon request from the corresponding author.

## Acknowledgements

This work was funded by the The Lustgarten Foundation (EMJ), NIH/NCI (P01CA247886 to EMJ; P30CA006973 and S10OD034407 to WJH), and the Dana & Albert R. Broccoli Charitable Foundation. SYM is supported by the ASCO & Conquer Cancer Foundation Young Investigator Award, the Bernice Garchik Pancreatic Cancer Fellowship, and the McGlothlin Fellows to Faculty Fund. We would also like to thank The Sidney Kimmel Comprehensive Cancer Center Flow Cytometry Technology Development Center, the Experimental and Computational Genomics Core, and Animal Resources.

---

**Supplementary Figure 1:**
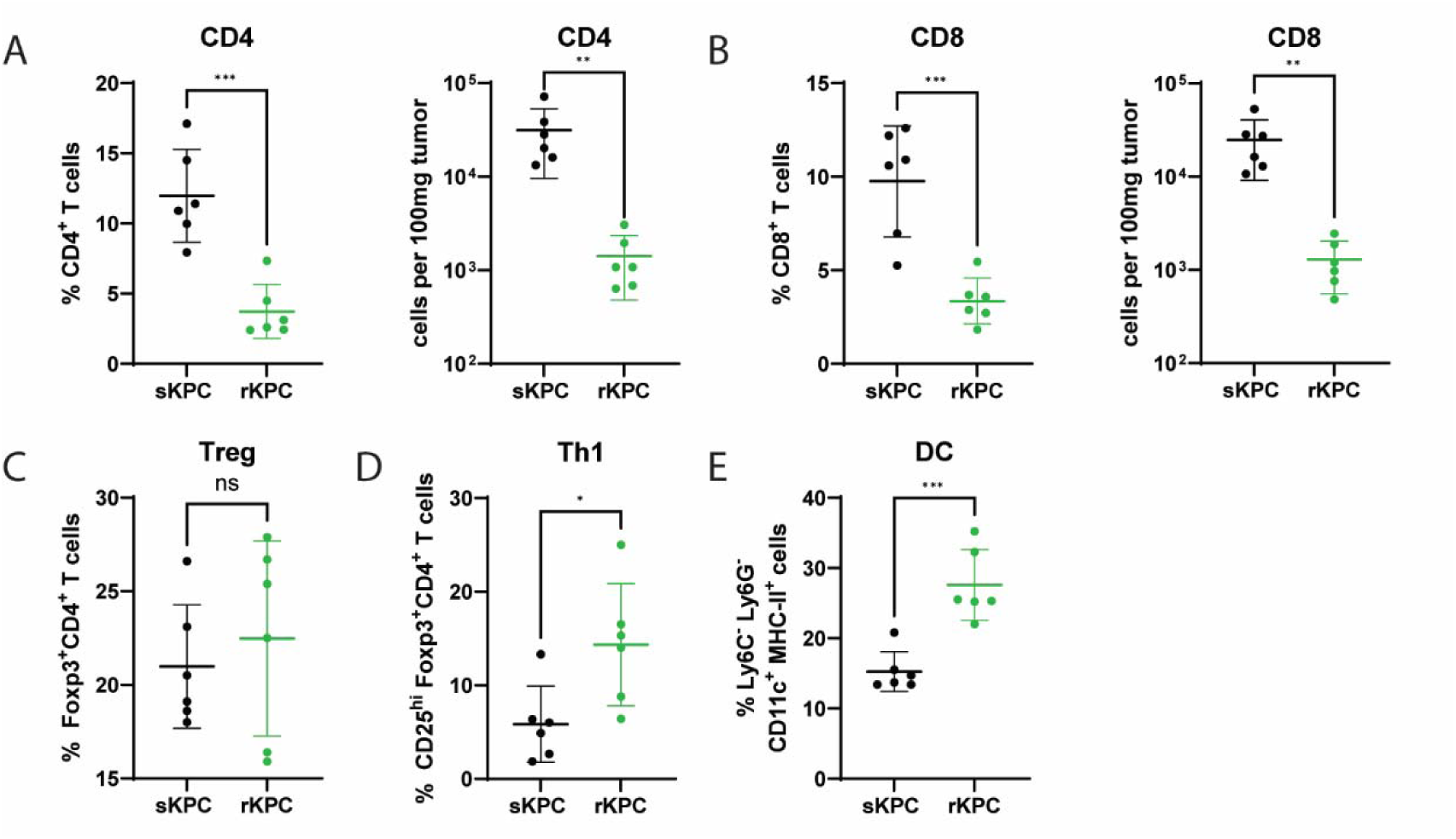
Characterization of immune cell infiltrates in sKPC and rKPC tumors. (A-B) Proportion of CD45^+^ cells (left) and absolute count per 100mg tumor (right) of CD4^+^ T cells (A) and CD8^+^ T cells (B). (C) Proportion of CD4^+^ T cells expressing Foxp3. (D) Proportion of CD4^+^ T cells expressing Tbet. (E) Proportion of DCs, defined as CD45^+^ F480^-^ Ly6C^-^ Ly6G^-^ cells expressing CD11c and MHC-II. Data shown is from one independent experiment, plotted as mean ± SD; n=6 mice per group.

**Supplementary Table 1:** Antibody list for imaging mass cytometry.

| Supplementary Table 1 |  |  |  |  |  |  |  |
| --- | --- | --- | --- | --- | --- | --- | --- |
| Mass | Metal | Antigen | Clone | Dilution Factor | Source | Catalogue | Custom |
| 113 | In | aSMA | 14A | 250 | Cell Signaling Technology | 69319SF | X |
| 145 | Nd | MHCII | M5/114.15.2 | 111.1 | Thermo Fisher | 14-5321-82 | X |
| 150 | Nd | Podoplanin | 8.1.1 | 250 | Biolegend | 916606 | X |
| 154 | Sm | Pan-Keratin | C11 | 250 | Cell Signaling Technology | 17171SF | X |
| 161 | Dy | CD3e | E4T1B | 250 | Cell Signaling Technology | 73484SF | X |
| 162 | Dy | CD4 | EPR19514 | 166.6 | Abcam | ab271945 | X |
| 164 | Dy | GFP | Polyclonal | 250 | Abcam | ab6673 | X |
| 165 | Ho | Foxp3 | FJK-16s | 250 | Standard BioTools | 3165024C |  |
| 191 | Ir | Cell-ID Intercalator Ir ("DNA1") | N/A | 400 | Standard BioTools | 201192B |  |

**Supplementary Table 2:**
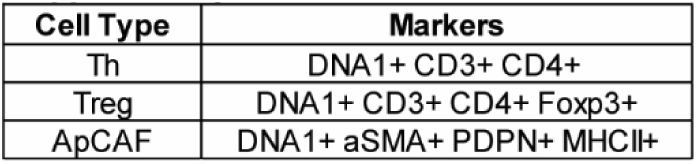
Phenotype definitions for proximity analysis.

| Supplementary Table 2 |  |
| --- | --- |
| Cell Type | Markers |
| Th | DNA1+ CD3+ CD4+ |
| Treg | DNA1+ CD3+ CD4+ Foxp3+ |
| ApCAF | DNA1+ aSMA+ PDPN+ MHCII+ |

**Supplementary Figure 2:**
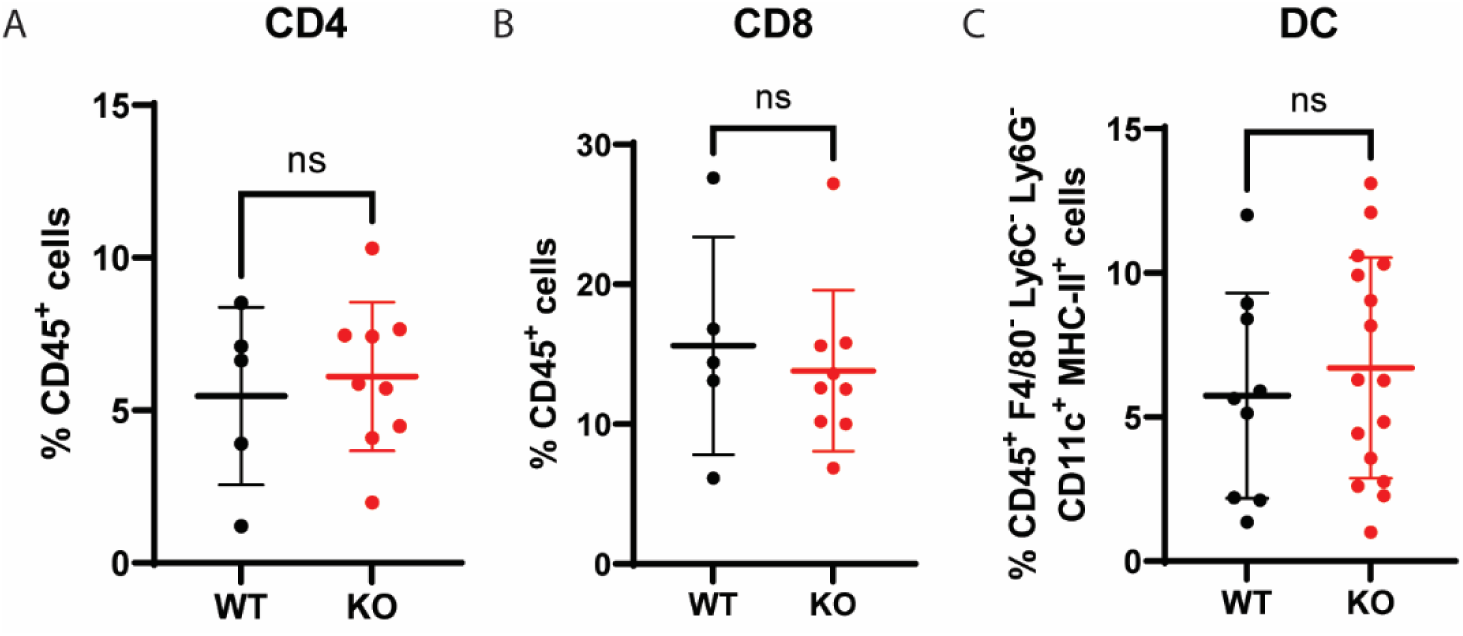
Characterization of immune cell infiltrates in apCAF KO and control mice. (A-B) Proportion of CD45^+^ cells of CD4^+^ T cells (A) and CD8^+^ T cells (B). (C) Proportion of DCs, defined as CD45^+^ F480^-^ Ly6C^-^ Ly6G^-^ cells expressing CD11c and MHC-II.

**Supplementary Figure 3:**
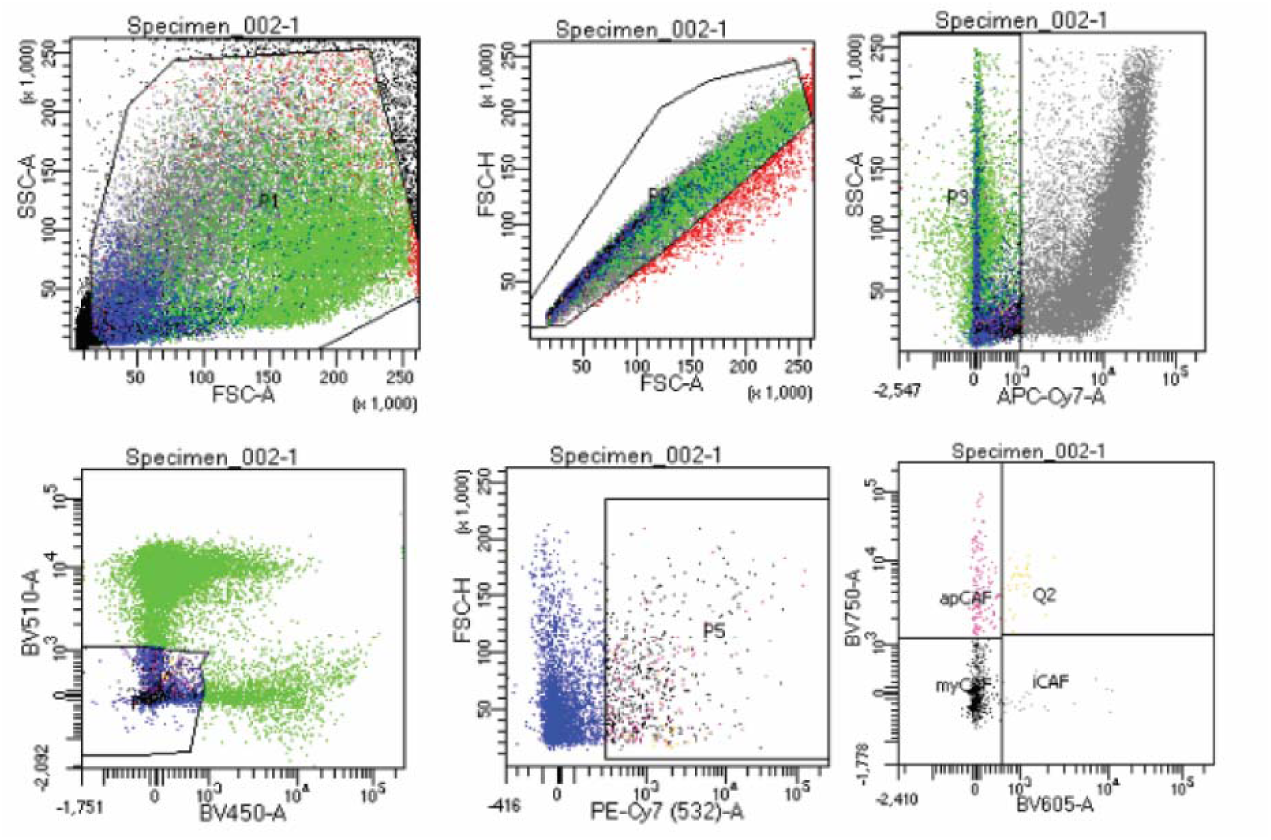
apCAF sorting strategy. apCAFs were sorted on a BD Fusion sorter. Cells were selected for size and singlets, followed by gating on cells negative for Zombie NIR live/dead stain. Cells negative for CD45 BV510 and dump (e-cadherin, EpCAM, CD31) on BV421 were gated, followed by gating on the PDPN PECy7 positive population to represent total CAFs. apCAFs were then defined as MHC-II BV785^+^ and Ly6C BV605^-^.

**Supplementary Figure 4:**
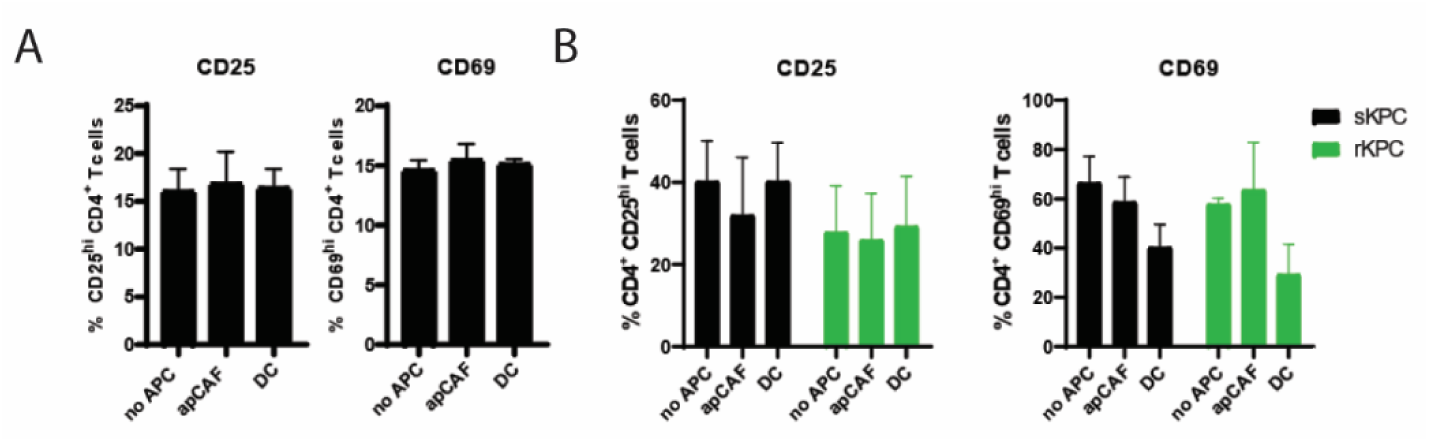
APCs without addition of cognate peptide do not activate CD4 T cells. **(A)** Sorted apCAFs and bead-isolated CD11c^+^ cells from sKPC tumors were co-cultured with tumor-infiltrating CD4^+^ T cells without addition of exogenous peptides. CD25 (left) and CD69 (right) expression on tumor-infiltrating CD4^+^ T cells. **(B)** Sorted apCAFs and bead-isolated CD11c^+^ cells from sKPC and rKPC tumors were pulsed with KRAS_G12D_ and co-cultured with tumor-infiltrating CD4^+^ T cells. CD25 (left) and CD69 (right) expression on tumor-infiltrating CD4^+^ T cells.

**Supplementary Figure 5:**
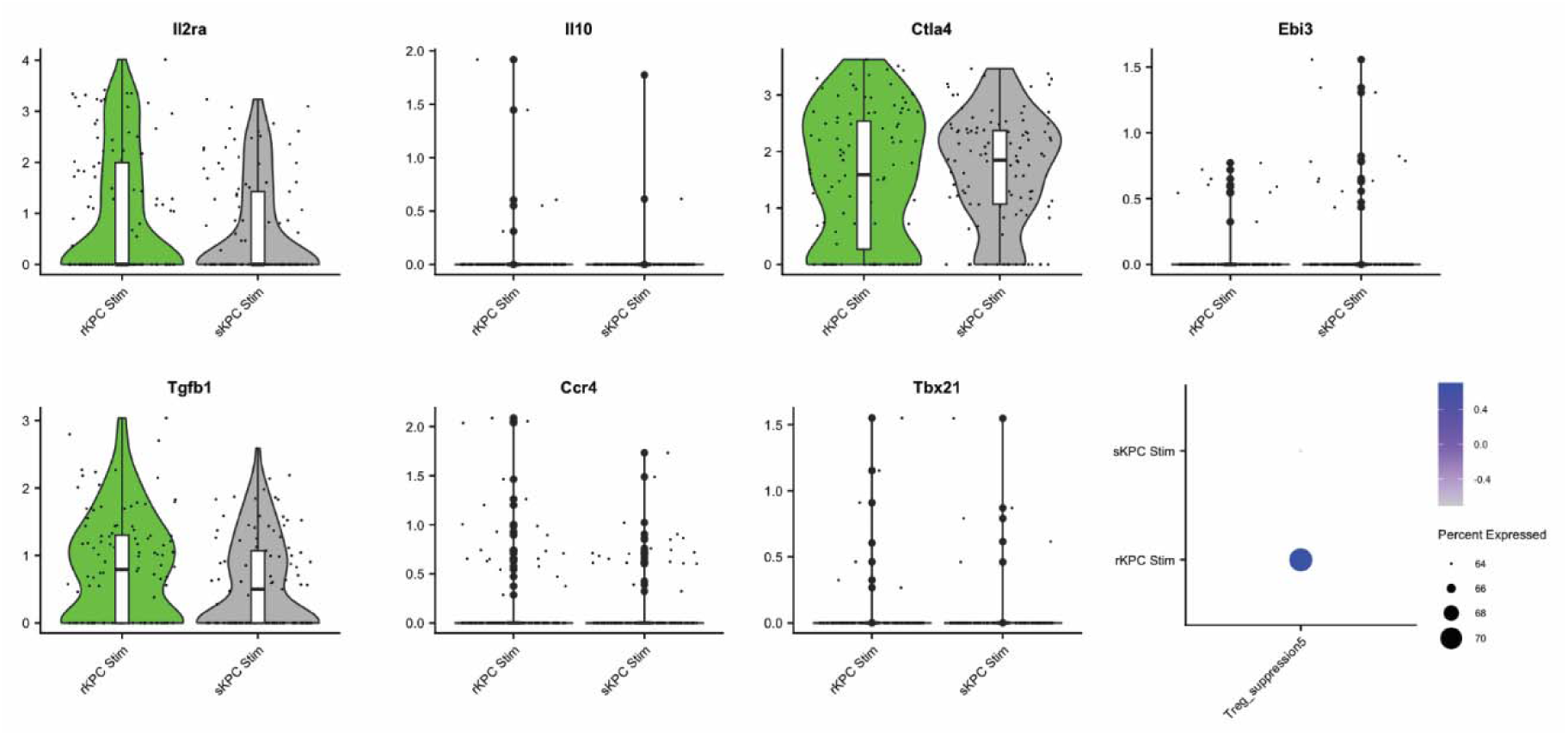
Expression of genes associated with Treg suppression. Gene expression of the individual genes included in the “Treg suppression” gene set between Tregs stimulated by sKPC apCAFs and rKPC apCAFs.

**Supplementary Figure 6:**
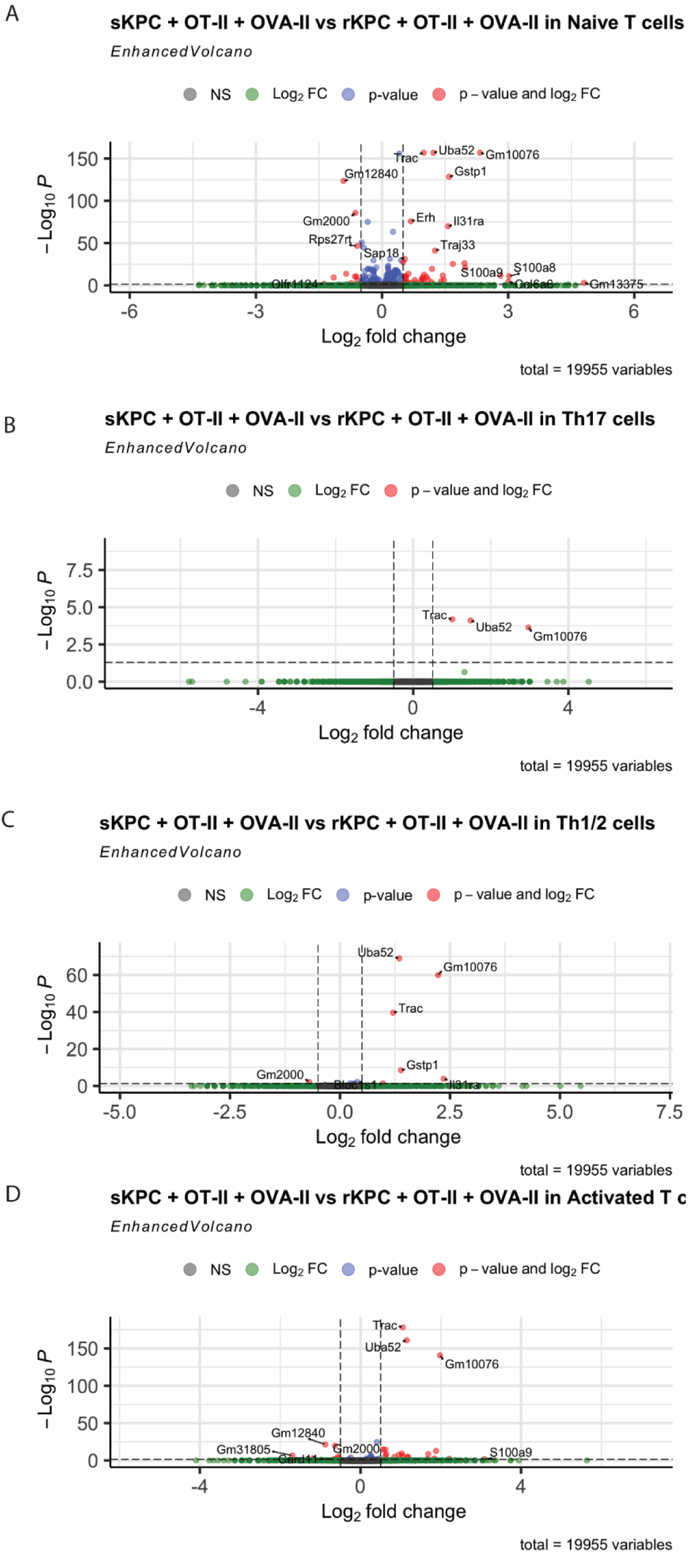
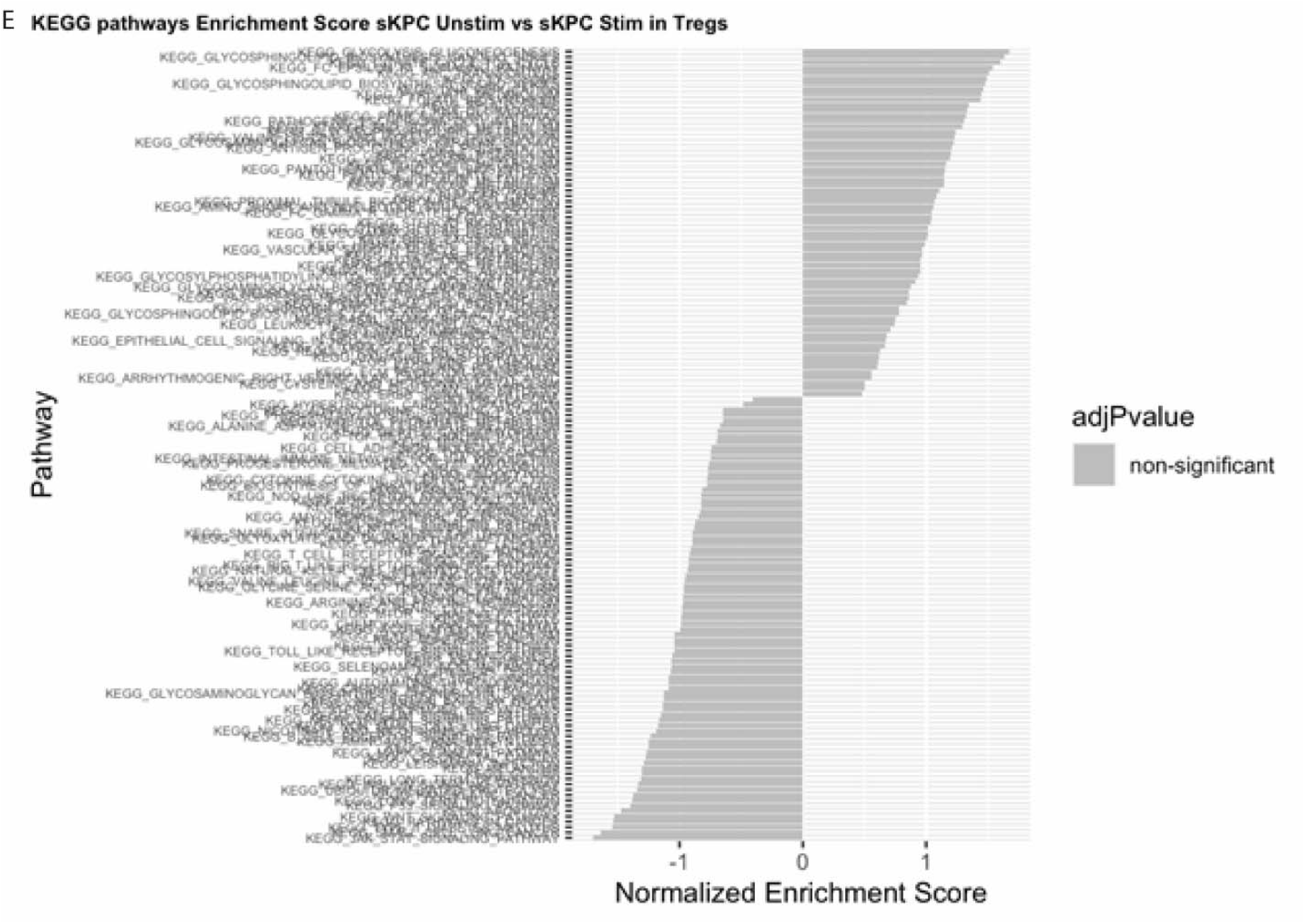
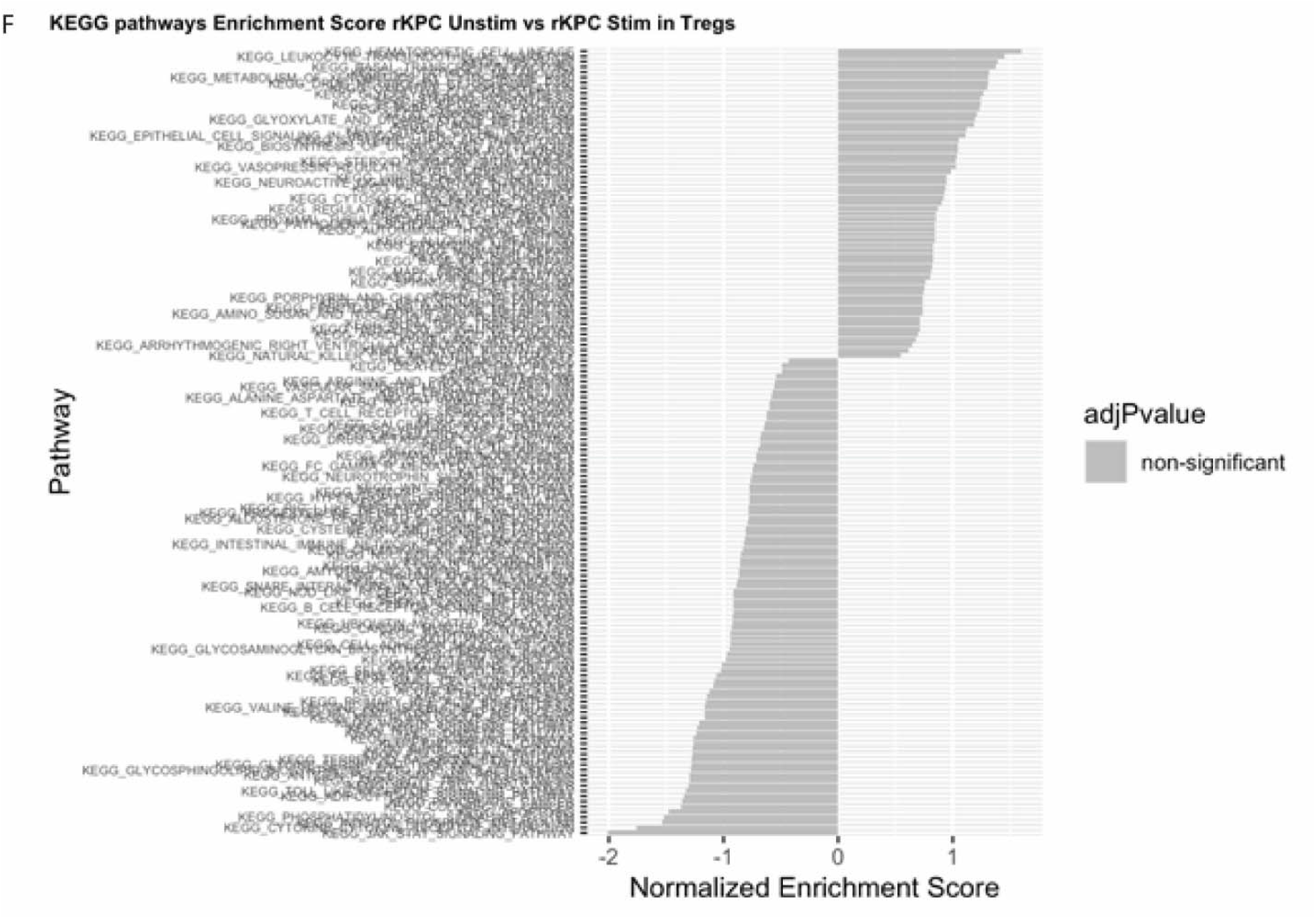
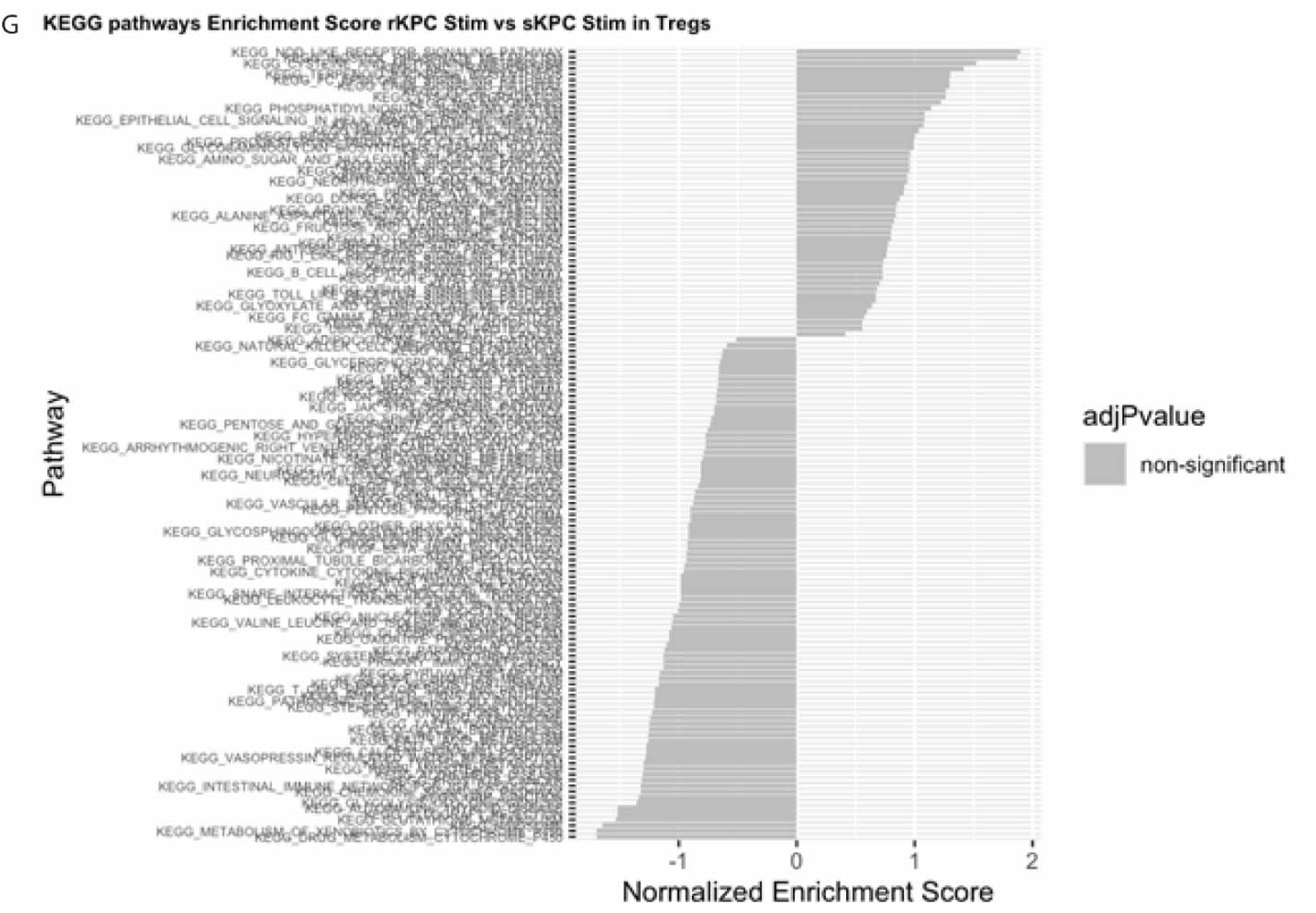
DEG and KEGG pathways upregulated in CD4^+^ T cell clusters. **(A-D)** Volcano plots showing differentially expressed genes in CD4^+^ T cells cocultured with pulsed sKPC apCAFs (right) versus pulsed rKPC apCAFs (left). **(A)** DEG of naive CD4^+^ T cells. **(B)** DEG of Th1/2 cells. **(C)** DEG of Th17 cells. **(D)** DEG of activated CD4^+^ T cells. **(E-G)** KEGG pathways for Tregs cocultured with sKPC apCAFs **(E)**, rKPC apCAFs **(F)**, and pulsed sKPC and rKPC apCAFs **(G)**.

**Supplementary Figure 7:**
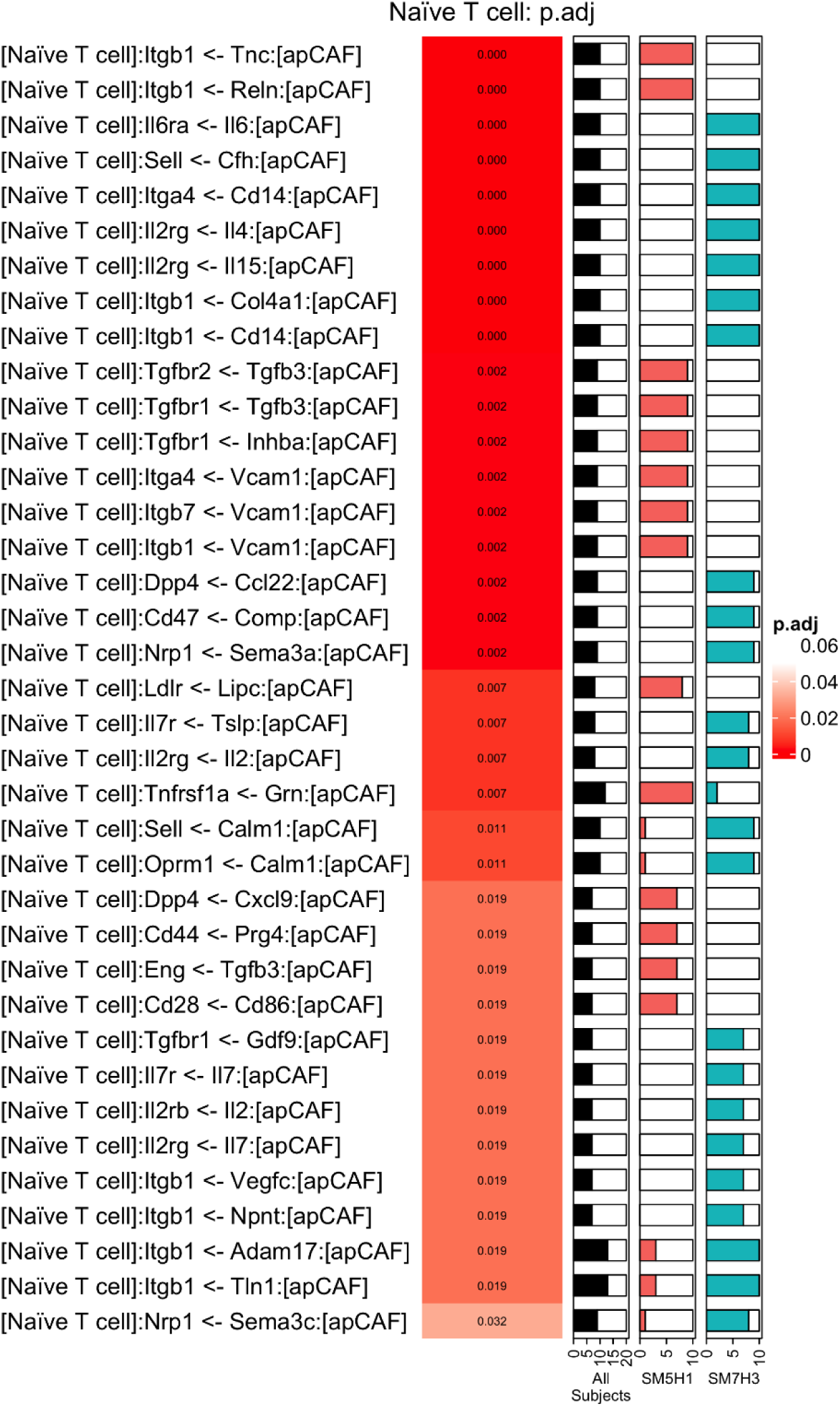
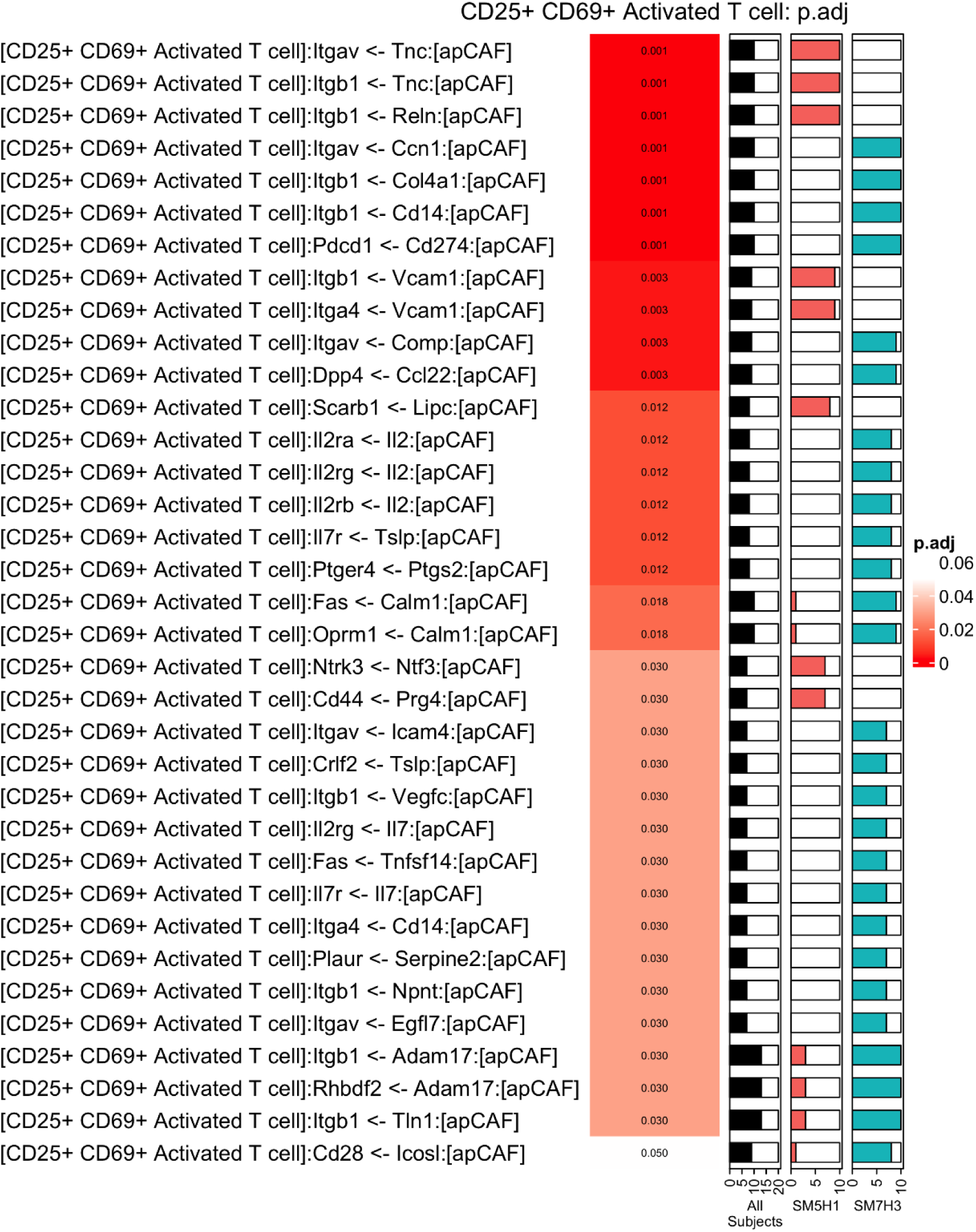
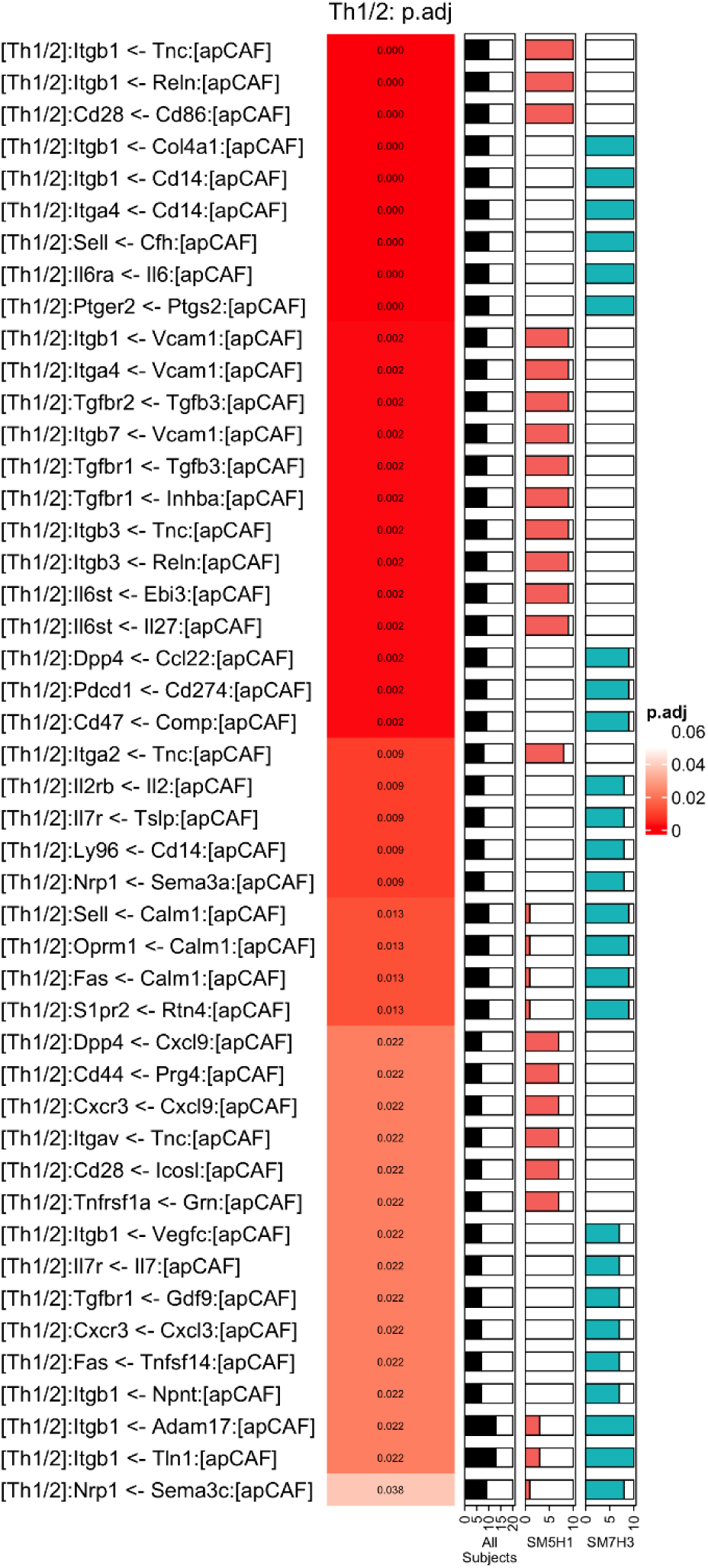
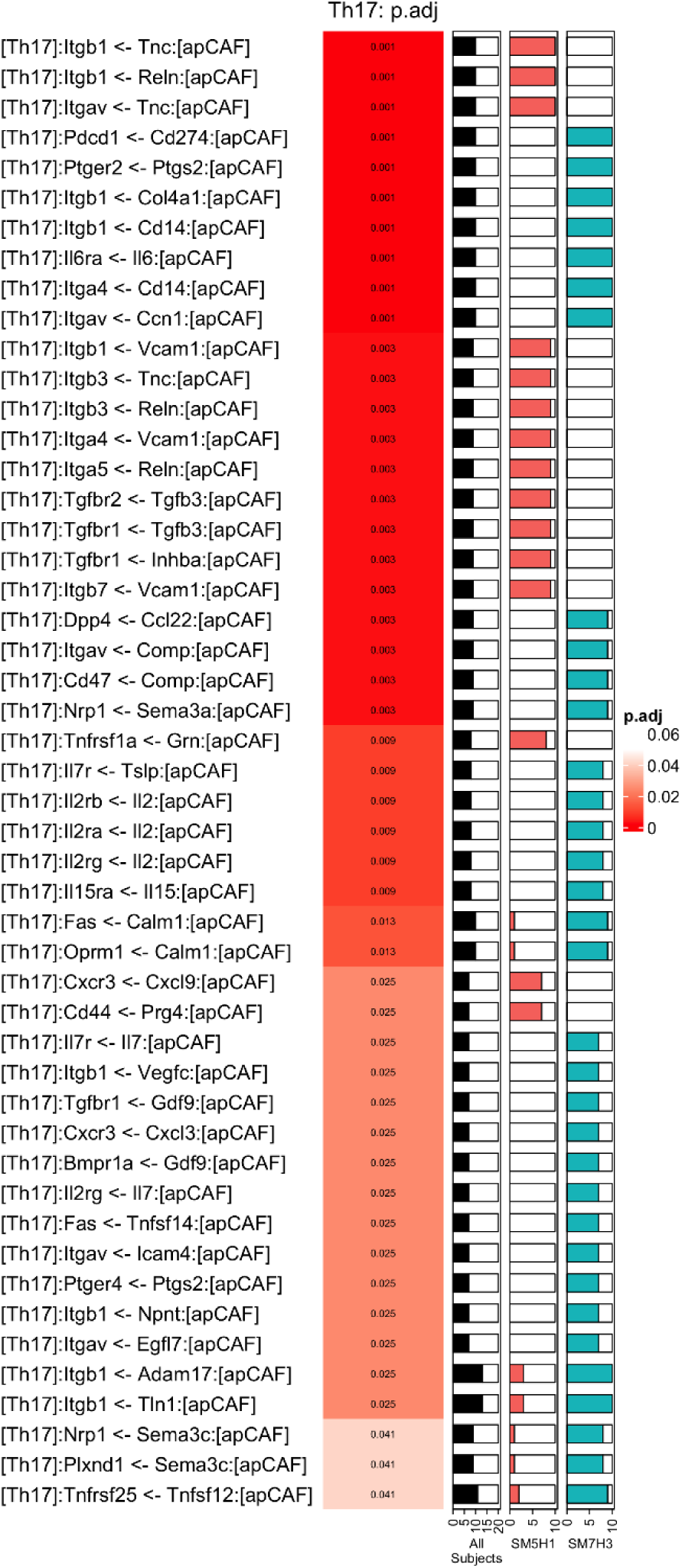
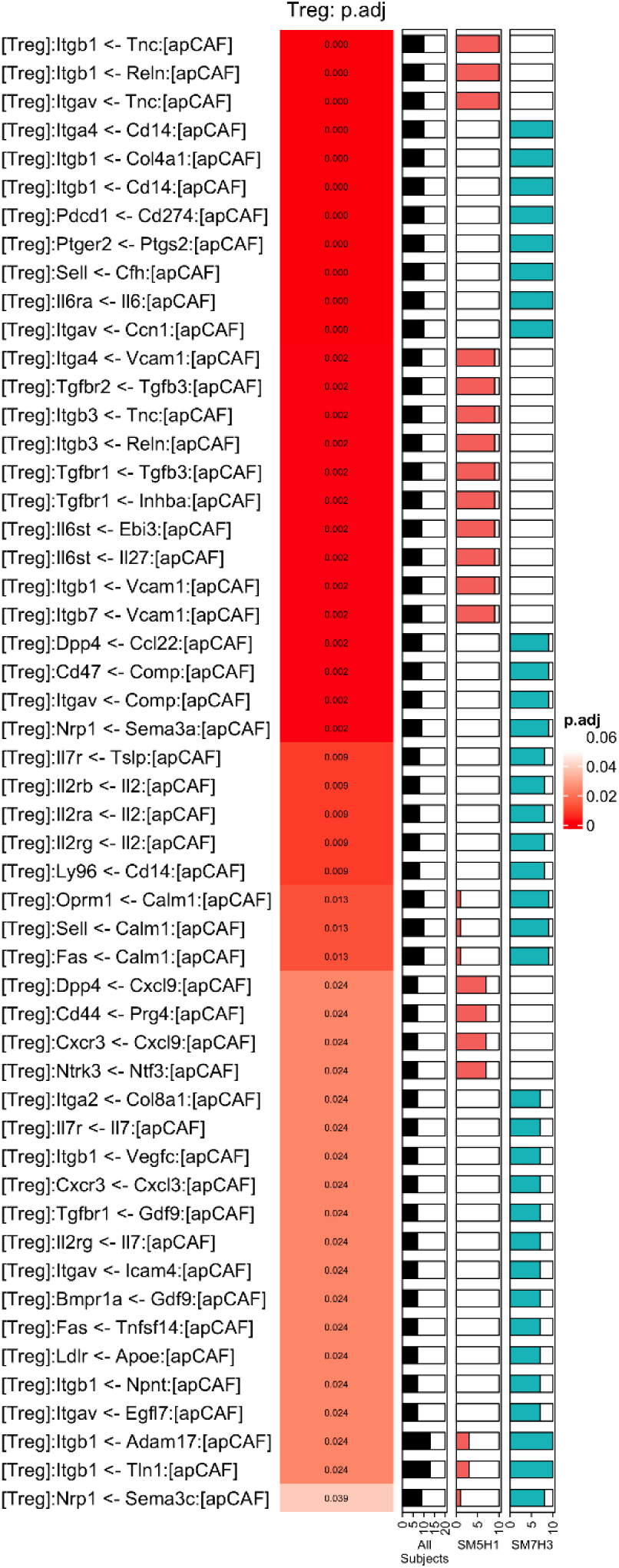
Cell to cell communication inference using dominoSignal. 10 bootstrapped samples of apCAFs and CD4^+^ T cells were generated from sKPC and rKPC experiments. Each bootstrapped sample was subjected to cell-cell communication inference using dominoSignal (v 1.1.0) and the dependence of inferred ligand-receptor signals from apCAFs to functional CD4 T cell subsets upon whether cells came from sKPC or rKPC tumors was tested by Fisher’s Exact test. Intercellular signaling interactions are named on a [‘receiver cell’]:’receptor’ <-‘ligand’:[‘sender cell’] basis. The left column is colored by the FDR-adjusted p-value of the Fisher’s exact test for occurrence of intercellular signals being inferred as active in each tumor type. The bars on the right show the proportion of all bootstrapped samples (grey), sKPC samples (black), and rKPC samples (green) where the signal was inferred as active by dominoSignal.

